# Cyanide-dependent control of terminal oxidase hybridization by *Pseudomonas aeruginosa* MpaR

**DOI:** 10.1101/2023.05.31.543164

**Authors:** Marina K. Smiley, Doran C. Sekaran, Alexa Price-Whelan, Lars E.P. Dietrich

**Affiliations:** Department of Biological Sciences, Columbia University, New York, NY 10025

## Abstract

*Pseudomonas aeruginosa* is a common, biofilm-forming pathogen that exhibits complex pathways of redox metabolism. It produces four different types of terminal oxidases for aerobic respiration, and for one of these–the *cbb*_3_-type terminal oxidases–it has the capacity to produce at least 16 isoforms encoded by partially redundant operons. It also produces small-molecule virulence factors that interact with the respiratory chain, including the poison cyanide. Previous studies had indicated a role for cyanide in activating expression of an “orphan” terminal oxidase subunit gene called *ccoN4* and that the product contributes to *P. aeruginosa* cyanide resistance, fitness in biofilms, and virulence–but the mechanisms underlying this process had not been elucidated. Here, we show that the regulatory protein MpaR, which is predicted to be a pyridoxal phosphate-binding transcription factor and is encoded just upstream of *ccoN4*, controls *ccoN4* expression in response to endogenous cyanide. Paradoxically, we find that cyanide production is required to support CcoN4’s contribution to respiration in biofilms. We identify a palindromic motif required for cyanide- and MpaR-dependent expression of *ccoN4* and co-expressed, adjacent loci. We also characterize the regulatory logic of this region of the chromosome. Finally, we identify residues in the putative cofactor-binding pocket of MpaR that are required for *ccoN4* expression. Together, our findings illustrate a novel scenario in which the respiratory toxin cyanide acts as a signal to control gene expression in a bacterium that produces the compound endogenously.

**IMPORTANCE:** Cyanide is an inhibitor of heme-copper oxidases, which are required for aerobic respiration in all eukaryotes and many prokaryotes. This fast-acting poison can arise from diverse sources, but mechanisms by which bacteria sense it are poorly understood. We investigated the regulatory response to cyanide in the pathogenic bacterium *Pseudomonas aeruginosa*, which produces cyanide as a virulence factor. Although *P. aeruginosa* has the capacity to produce a cyanide-resistant oxidase, it relies primarily on heme-copper oxidases and even makes additional heme-copper oxidase proteins specifically under cyanide-producing conditions. We found that the protein MpaR controls expression of cyanide-inducible genes in *P. aeruginosa* and elucidated the molecular details of this regulation. MpaR contains a DNA-binding domain and a domain predicted to bind pyridoxal phosphate (vitamin B6), a compound that is known to react spontaneously with cyanide. These observations provide insight into the understudied phenomenon of cyanide-dependent regulation of gene expression in bacteria.

## INTRODUCTION

Biofilms are crowded structures containing cells and secreted polymers that form an adherent matrix. As a biofilm grows, internal chemical gradients become established because (i) cells near the biofilm edge consume substrates such as O_2_ before these resources can travel through the structure and (ii) cells in separate biofilm subregions release distinct products. Cells in microniches within biofilms use different tactics to survive, thus decreasing the efficacy of single-target treatments (1–5).

The opportunistic pathogen *Pseudomonas aeruginosa* is known for its respiratory versatility and its production of small molecules that interact with the respiratory chain (6–9). Depending on the growth conditions, it can synthesize four major types of terminal oxidases, i.e. enzyme complexes that catalyze the final step of aerobic respiration, including high-affinity *cbb*_3_-type oxidases (“Ccos”). The production of small molecules that interact with the respiratory chain is generally controlled by quorum-sensing pathways and/or regulators that respond to low-oxygen conditions (7, 10, 11). Among these compounds are the *P. aeruginosa* phenazines, a class of redox-cycling compounds that have been implicated in anaerobic survival (12, 13) and in electron transfer by the respiratory chain (9, 14). Together, the terminal oxidases and phenazines provide extensive and varied potential routes of metabolic electron flow for *P. aeruginosa* cells in biofilms. Adding to this complexity is the fact that *P. aeruginosa* has the capacity to produce cyanide, a potent inhibitor of heme copper oxidases including *cbb*_3_-type oxidases (15–18). These observations raise questions about how *P. aeruginosa* mediates electron flow when O_2_ limitation and cyanide production threaten respiratory function.

The *P. aeruginosa* genome contains multiple, differentially regulated genetic loci that produce full or partial Cco complexes, with functionally interchangeable subunits (19) (**Figure 1A,B**). One of genes, *ccoN4*, encodes a so-called “orphan” Cco subunit that contributes to WT biofilm formation and competitive fitness in biofilms (14). *ccoN4* is part of a cluster that is strongly induced by cyanide (20). At first, it seems odd that cyanide induces *ccoN4* expression because cyanide is known for its specific inhibition of oxidases like the Ccos (21). Nevertheless, studies have shown that *ccoN4* contributes to resistance when exogenous cyanide is added to liquid cultures (19, 20). Though *P. aeruginosa* can produce a cyanide-insensitive oxidase (CIO) that does not contain a heme-copper center, a drawback to the use of this oxidase is that it has a low affinity for O_2_ relative to the Cco complexes. Moreover, viability studies have shown that CIO is not the sole contributor to *P. aeruginosa* cyanide resistance (19, 22). These observations suggest that the CcoN4 subunit is functional in the presence of cyanide and that its use may be preferred when high O_2_ affinity is beneficial such as in hypoxic biofilm settings.

**Figure 1.**
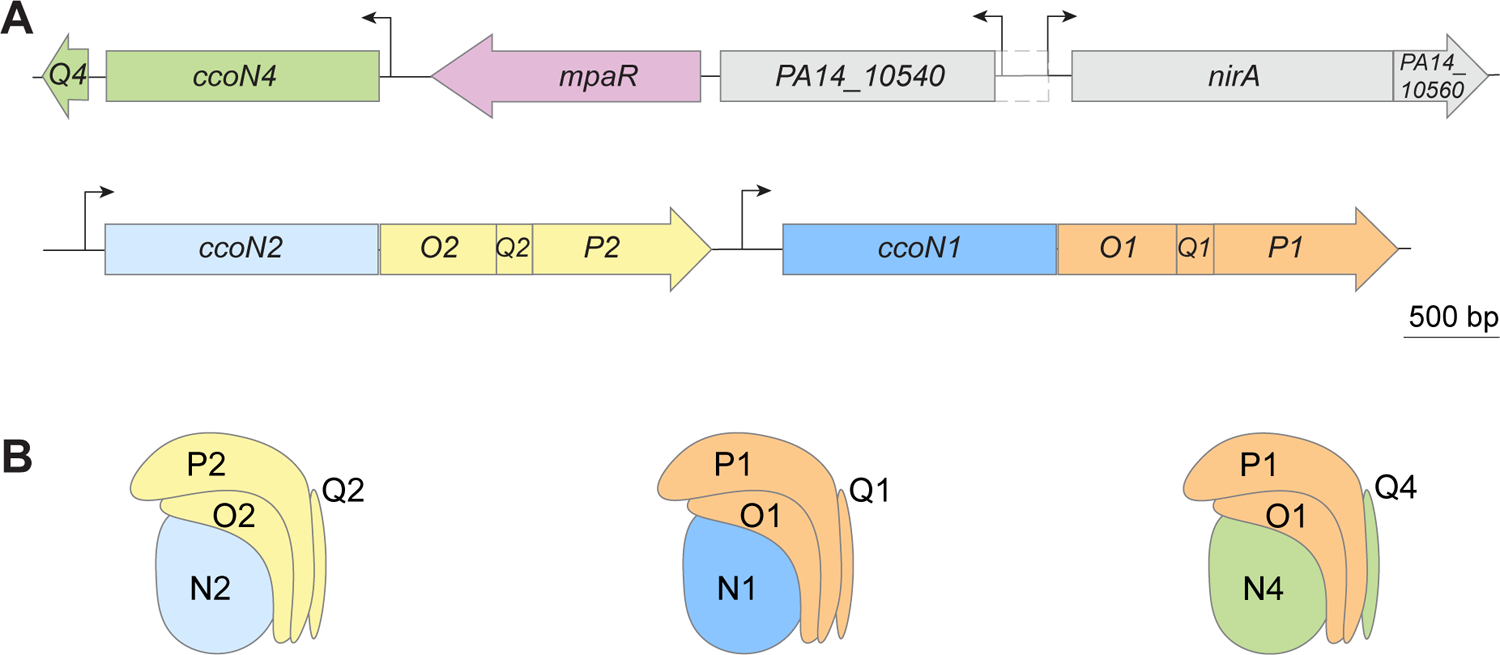
Genomic arrangement of *cco* genes. (**A**) Arrangement of *mpaR* and *cco* genes on the *P. aeruginosa* PA14 chromosome. The locations of transcription start sites were determined using data in Wurtzel et al. 2012 (35). The depicted size of the *PA14_10540* ORF is based on the TSS indicated by RNAseq data; a larger ORF (area with dashed outline) is predicted by The Pseudomonas Genome Database and BioCyc. (**B**) Cartoon representations of assembled *cbb*_3_-type terminal oxidase complexes, based on the Cco structure from *P. stutzeri* (PDB: 3mk7) (88) and including a hybrid isoform that combines Cco subunits encoded by different operons.

*ccoN4* lies adjacent to *mpaR* (*PA14_10530*), which codes for a GntR family transcription factor. In this study, we investigated the role of MpaR in *ccoN4* and *mpaR* expression and its responsiveness to cyanide. Our findings support a model for the role of this regulatory protein in cyanide resistance across depth in *P. aeruginosa* biofilms.

## RESULTS

### CcoN4 contributes to TTC reduction in a cyanide-dependent manner

Among the various types of terminal oxidases produced by *P. aeruginosa*, the *cbb*_3_ type is most important for growth under standard laboratory conditions and appears to be the only one that is produced constitutively (23–26). A minimal functional *cbb*_3_-type complex comprises two subunits, CcoN and CcoO; two additional and nonessential subunits, CcoP and CcoQ, have been implicated in assembly or electron transfer (27–29). Although *cco* genes (encoding *cbb*_3_ type oxidase subunits) are found in diverse bacteria, the *P. aeruginosa* genome is unusual in that it contains two complete and two partial *cco* operons (14, 19). The *cco1* and *cco2* operons lie adjacent to each other and are each sufficient to produce a full Cco complex (**Figure 1A,B**); the *ccoN3Q3* and *ccoN4Q4* operons are found at independent locations in the genome and code for orphan subunits that can replace their corresponding subunits in Cco1 or Cco2 to produce hybrid complexes. A study conducted in *P. aeruginosa* PAO1 tested 16 possible isoforms of Cco, containing subunits encoded by different operons in distinct combinations, and demonstrated that they were all sufficient to support aerobic growth similar to that seen for the parent strain (19). However, several lines of evidence point to unique physiological roles for the CcoN4 subunit. Our prior work in *P. aeruginosa* PA14 has implicated *ccoN4* in (i) phenazine reduction and fitness during biofilm growth, and (ii) virulence in a *C. elegans* slow-kill assay (14). In addition, work by others has indicated that *ccoN4* contributes specifically to resistance to exogenous cyanide (19, 20). These observations motivated us to test the contribution of CcoN4 to respiration in the presence of endogenously produced cyanide.

To ask whether endogenous cyanide affects respiration via specific CcoN subunits, we used a tetrazolium chloride (TTC)-based assay in which the colorless dye changes to red when reduced by cytochrome c oxidase-dependent respiration (30) (**Figure 2A**). We examined TTC reduction in *P. aeruginosa* PA14 biofilms of single and combinatorial mutants lacking *ccoN* genes and/or the *hcnABC* operon (“*hcn*”), the latter of which is required for cyanide synthesis. We found that CcoN4 contributes to dye reduction in this assay, particularly in the Δ*N1*Δ*N2* background, as described previously (14). We also found that this contribution is dependent on the presence of *hcnABC* (**Figure 2A**). This result is consistent with the prior observation, made in *P. aeruginosa* strain PAO1, that endogenous cyanide induces *ccoN4Q4* expression (20). Paradoxically, the role of cyanide in *ccoN4* induction appears to create a scenario where a respiratory toxin is necessary for substantial respiration in the Δ*N1*Δ*N2* background.

**Figure 2.**
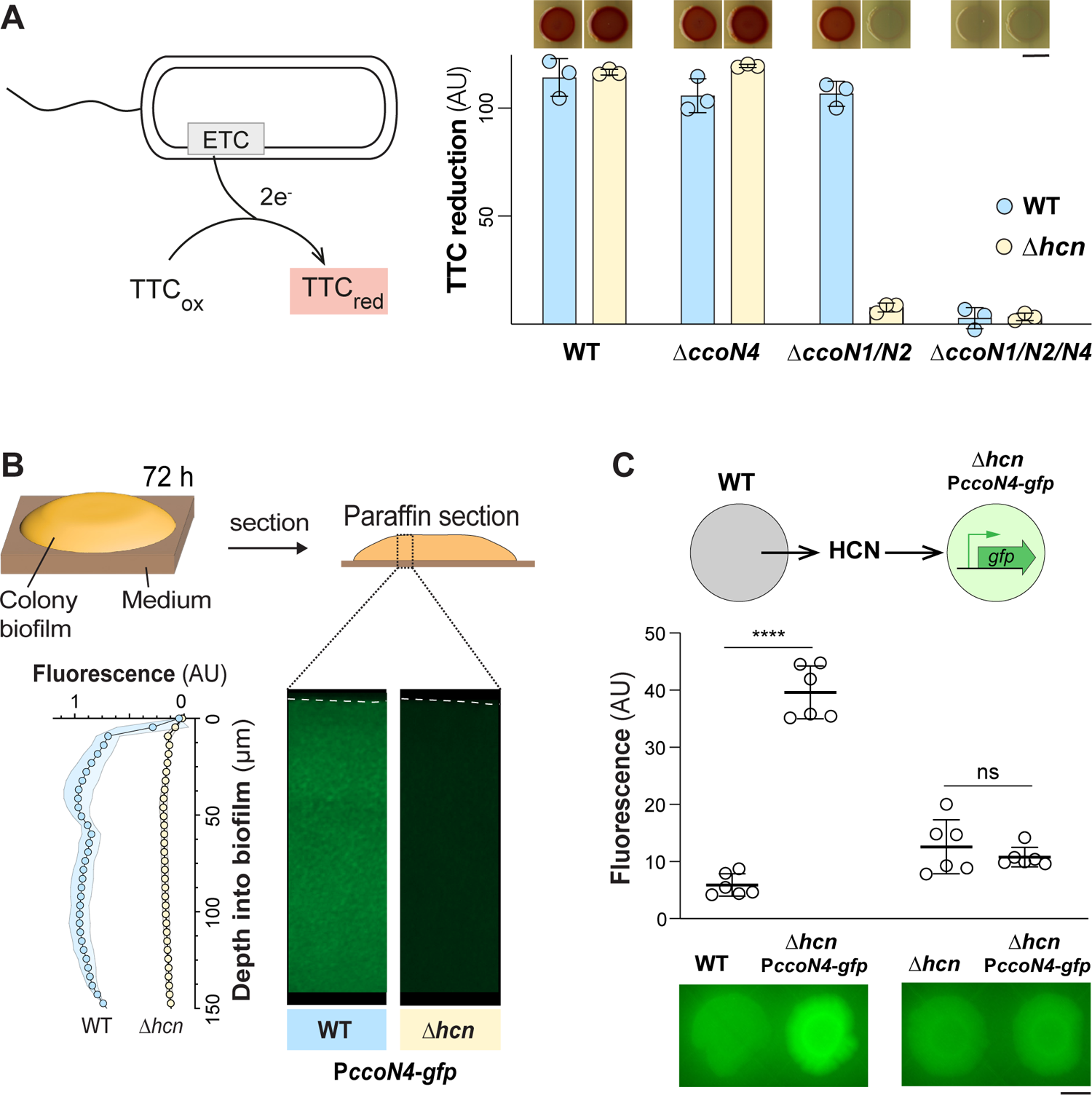
*ccoN4* contributes to respiration and is induced by endogenous cyanide. (**A**) Left: TTC reduction by the electron transport chain (ETC) produces a red compound. Right: TTC reduction by colony biofilms of the indicated strains, quantified as the average saturation in the red hue for all pixels in the colony area. Data points represent biological replicates and error bars indicate standard deviation. Scale bar is 5 mm. (**B**) Top: Colony biofilms were grown for 3 days before harvesting and preparation of thin sections. Bottom left: Activity of the P*ccoN4-gfp* reporter construct across depth in colony biofilms. Bottom right: Representative thin-section images of the indicated strains, each containing the P*ccoN4-gfp* reporter construct. Dotted lines indicate the top of each biofilm. Experiments were performed with biological triplicates and shading indicates standard deviation. (**C**) Top: Assay for HCN-dependent induction at a distance. Middle: Top-down quantification of fluorescence for adjacent biofilms grown in the HCN-dependent induction assay. Error bars represent the standard deviation of biological triplicates. Biofilms were grown for 3 days on 1% tryptone 1% agar before scanning. Scale bar is 5 mm.

### Endogenous cyanide boosts transcription of *ccoN4* in *P. aeruginosa* PA14 biofilms

Cyanide-dependent control of *ccoN4Q4* expression has previously been characterized in *P. aeruginosa* PAO1 liquid cultures (19, 20). However, cyanide synthesis is regulated in response to cell density and studies comparing PAO1 and PA14 have described strain-dependent differences in quorum-sensing controlled processes (31, 32). Furthermore, the condition-dependence of cyanide production (10, 33) could affect its regulation in chemically heterogeneous biofilm subzones (10, 33). We therefore sought to directly examine the effect of endogenously produced cyanide on activity of the *ccoN4* promoter in PA14 liquid cultures and biofilms. We created a reporter construct in which *ccoN4*’s putative promoter region drives expression of *gfp* and moved it into a neutral chromosomal site in the WT and a Δ*hcn* mutant (34). Fluorescent growth curves show that the *ccoN4* transcription is induced at the onset of stationary phase and that total expression levels are lower in the Δ*hcn* background (**Figure S1**). This result is in line with measurements of P*ccoN4*-*lacZ* reporter activity made by Hirai et al., though the effect of *hcn* gene deletion in their study was more pronounced (19). We conclude that cyanide boosts but is not entirely required for transcription of *ccoN4* in PA14 liquid cultures.

To examine the role of cyanide in *ccoN4* expression in biofilms, we grew macrocolony biofilms for 3 days on 1% tryptone 1% agar medium, embedded them in paraffin, and prepared thin sections for microscopic imaging. We focused our analysis on the top 150 µm of the biofilm, which includes the transition from oxic to anoxic subzones (14), because the syntheses of cyanide and some terminal oxidases are oxygen-regulated (10, 24, 26). In the WT background, the *ccoN4* transcriptional reporter showed high levels of expression across most of the biofilm depth, with a reproducible peak in expression at a depth of 40-50 µm (**Figure 2B**). *ccoN4* expression across depth was greatly reduced in the cyanide-null background. Finally, because cyanide is a volatile compound, cyanide-producing strains should be able to chemically complement cyanide-null mutants when they are grown in proximity. To test this, we grew WT and Δ*hcn* biofilms near, but not touching, Δ*hcn* P*ccoN4-gfp* biofilms on an agar plate. We found that incubation of cyanide-null biofilms near WT (cyanide-producing) biofilms was sufficient to activate *ccoN4* expression (**Figure 2C, D**). In sum, PA14 planktonic cultures and biofilms both show enhancement of *ccoN4* expression by endogenous cyanide, with biofilms showing a greater enhancement than planktonic cultures due to lower basal expression.

### MpaR is required for *ccoN4* expression

In *P. aeruginosa* strain PAO1, high induction by cyanide was observed not just for the *ccoN4Q4* operon (ORFS PA4133-4134) but also the four genes upstream, which are ORFS PA4129-PA4132 (20). The chromosomal arrangement of the six corresponding genes in strain PA14 is shown in **Figure 1A**. Transcriptional analyses indicate that the gene pairs *nirA*-*PA14_10560* and *PA14_10540-mpaR* are each co-transcribed (35). *nirA* codes for a nitrite reductase (36) and *PA14_10540* codes for an ortholog of CcoG, a cupric reductase involved in the assembly of *cbb*_3_-type oxidases (37). We were intrigued by the presence of *mpaR*, coding for a GntR-family transcriptional regulator, directly adjacent to *ccoN4* because transcription factors are often encoded upstream of (one of) their targets. A recent RNAseq analysis using a *P. aeruginosa* PAO1 Δ*mpaR* mutant and a 2-fold cutoff showed that MpaR affects the expression of 275 genes and highlighted its integration into *P. aeruginosa*’s quorum-sensing network (**Figure S2**) (38).

The transcripts for ORFs PA4129-PA4134 (which includes *ccoN4Q4*) were among those that showed the largest decreases in abundance in Δ*mpaR* when compared to the WT. To directly test whether MpaR is required for *ccoN4* expression in strain PA14, we created a markerless *mpaR* deletion (Δ*mpaR*) and inserted the P*ccoN4-gfp* reporter construct at a neutral chromosomal site. To test whether *mpaR* affects the expression of other *cco* genes, we also deleted *mpaR* in strains containing *gfp* reporter constructs for P*ccoN1* and P*ccoN2*. Finally, to test whether Anr, an O_2_-sensitive transcription factor required for *cco2* expression (26) is important for P*ccoN4* activity, we moved the P*ccoN4*-*gfp* construct into the Δ*anr* background. We used these reporter strains to assess promoter activity during growth in liquid cultures and across depth in biofilms.

We found that planktonic growth of the Δ*mpaR* strain is altered, relative to the WT, at the transition from exponential to stationary phase (**Figure 3A**). In addition to genes that enhance cyanide resistance, the MpaR regulon includes genes that code for (i) primary metabolic and respiratory enzymes, (ii) proteins that contribute to cell density-dependent processes (i.e., quorum sensing), and (iii) sigma factors, all of which may influence exponential-to-stationary phase growth kinetics. Examining reporter fluorescence in liquid cultures, we further found that MpaR is required for expression of *ccoN4* while its loss has minimal effect on *cco1* expression and only a partial effect on the expression of *cco2*, which is a well established target of Anr (24, 26, 39) (**Figures 3A**). Complementing the *mpaR* deletion restored *ccoN4* expression to WT levels (**Figure S3**). These observations are consistent with RNAseq data from Wang et al. 2020, in which the *ccoN4* and *ccoP2* transcripts were ∼98-fold and ∼2-fold, respectively, less abundant in a Δ*mpaR* mutant compared to its parent strain, *P. aeruginosa* PAO1 (38).

**Figure 3.**
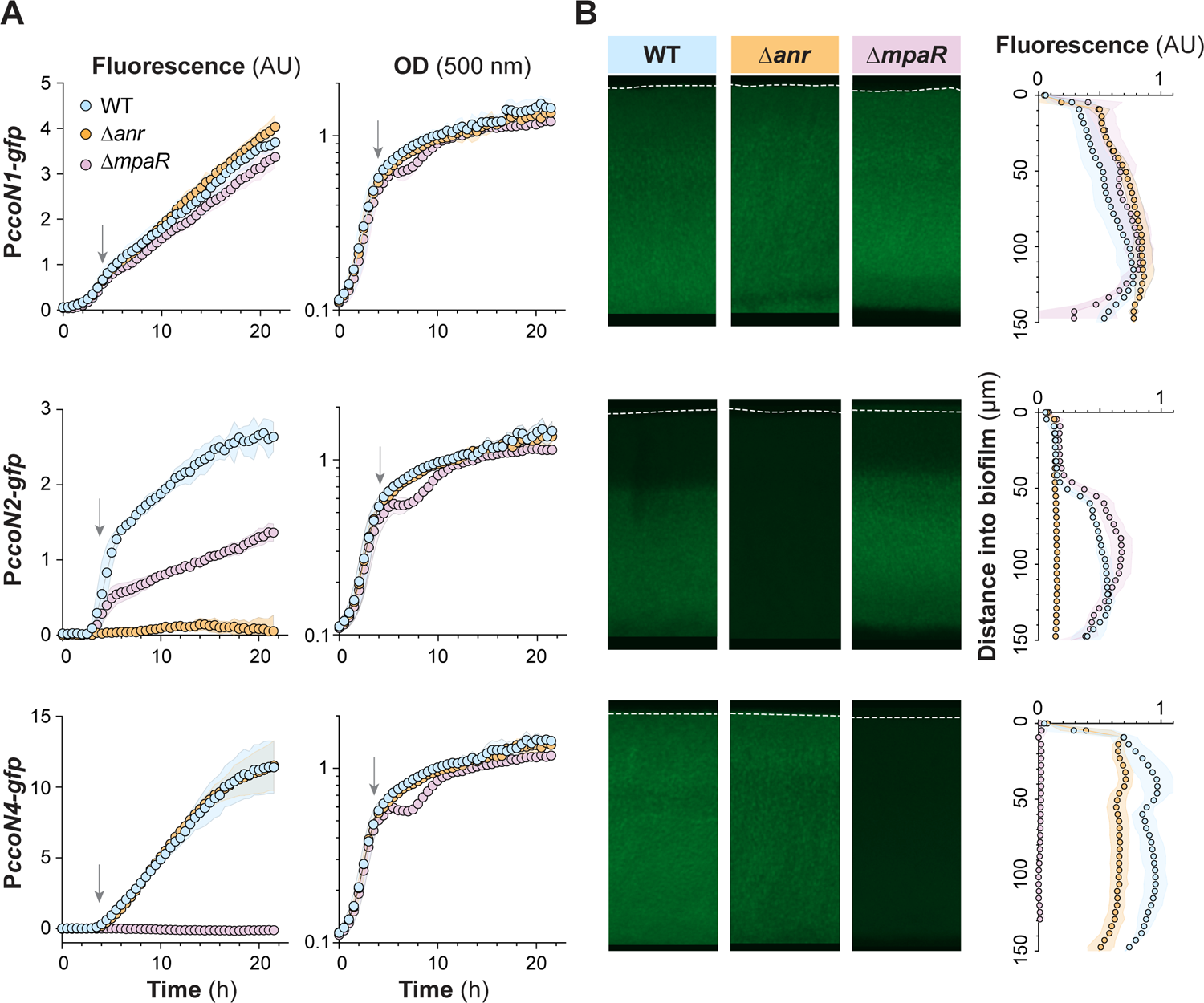
*P. aeruginosa* Anr and MpaR control *cco* gene expression in liquid cultures and biofilms. (**A**) Activities of the *ccoN1*, *ccoN2*, *ccoN4* promoters during growth in liquid culture (1% tryptone) reported as GFP fluorescence. Data represent the mean of six biological replicates with shading indicating standard deviation. Arrow represents the onset of stationary phase. (**B**) Thin-sections of biofilms grown for 3 days. The quantification on the right shows the mean of biological triplicates and shading indicates standard deviation.

Interestingly, in contrast to liquid culture, the loss of *mpaR* showed little to no effect on *cco2* expression in biofilms (**Figure 3**). Additionally, while we observed that deletion of *anr* does not affect *ccoN4* expression in planktonic growth, it has a small downregulatory effect during biofilm growth which may arise from the fact that Anr promotes cyanogenesis (10). In both modes of growth, reporter data indicated that neither Anr nor MpaR contribute to *cco1* expression, which is in line with prior studies (24). Finally, having observed that *ccoN4* contributes to respiration in biofilms and that MpaR regulates this gene, we tested whether respiratory activity was altered in Δ*mpaR* biofilms relative to those formed by the WT. We found that *mpaR* was required for WT levels of TTC reduction specifically in the cyanide-producing background, consistent with its role in inducing *ccoN4* (**Figure S4**).

### Identification of a palindromic motif required for *ccoN4* expression

We next sought to characterize features of the *ccoN4* promoter that underpin MpaR-dependent expression. According to a genome-wide analysis carried out by Wurtzel et al., the transcription start site (TSS) for *ccoN4* is 69 bp upstream of its start codon (**Figure 4A**) (35). To identify regions of the *ccoN4* promoter that are required for MpaR-dependent expression, we created a series of reporter constructs in which *ccoN4-*promoter sequences of varying length drove *gfp* expression. We found that shortening the promoter region from 263 to 138 bp decreased, but did not abrogate, *gfp* expression (**Figure 4B**), indicating the involvement of additional factors that promote *ccoN4* expression (in this context a GC-rich sequence at positions 147 to 138 bp upstream of the start codon is noteworthy (38)). However, further shortening the promoter length from 138 to 129 bp fully prevented expression of our *gfp* reporter (**Figure 4B**) indicating that the 138-129-bp sequence is required for *ccoN4* expression. Intriguingly, this stretch contains a short palindromic motif (ATCTGAT), which also appears in the promoter regions of the adjacent genes *PA14_10540* and *nirA*—the two other targets in the vicinity that may also be under the control of MpaR (**Figure 4C**) (38). We shuffled these seven base pairs (ATCTGAT to GATATCA) in the full-length promoter construct and found that they are required for gene expression (**Figure 4B**).

**Figure 4.**
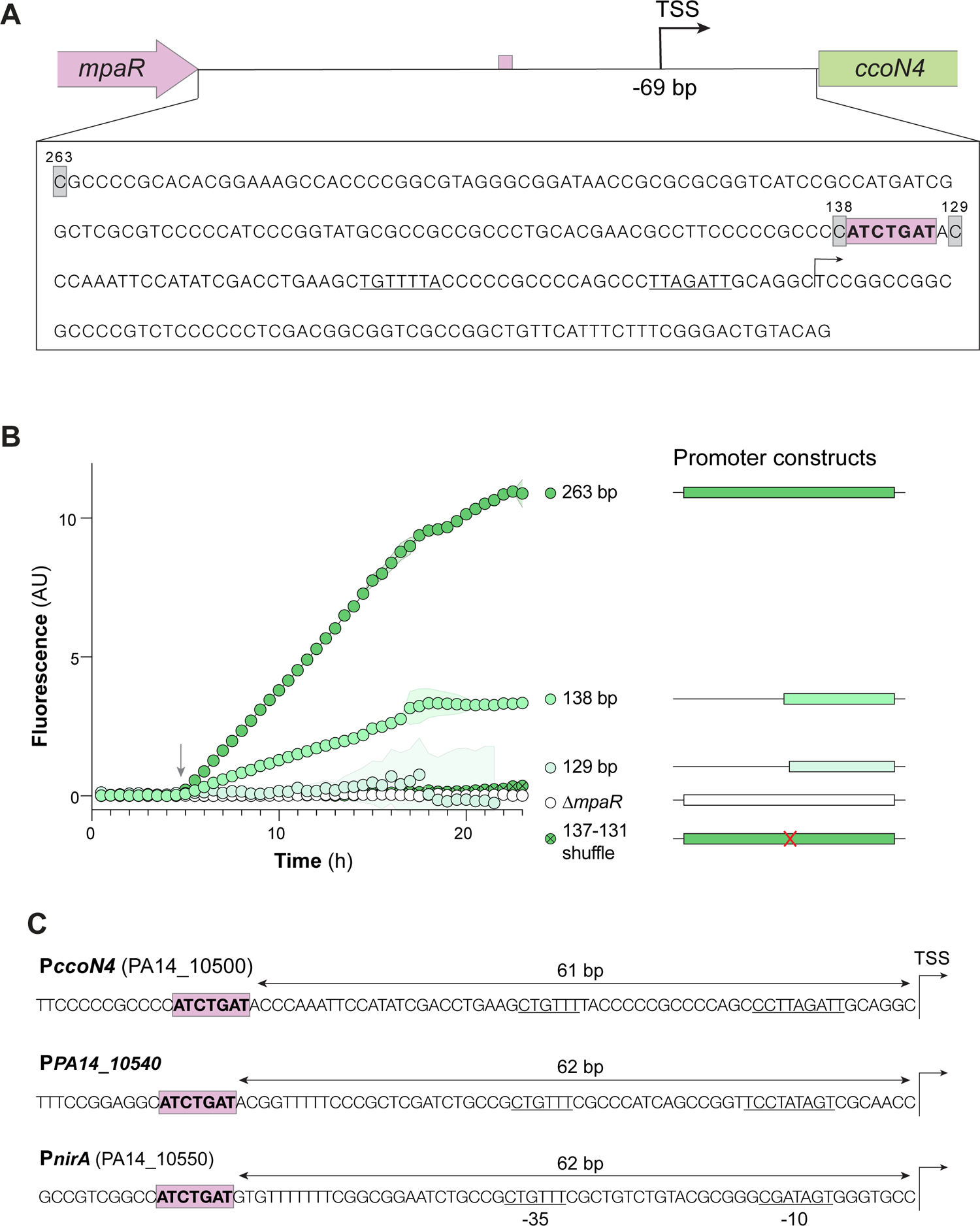
Mutational analysis of the *ccoN4* promoter region reveals a putative MpaR binding site. (**A**) Top: Schematic showing genomic orientation and transcription start site of the *mpaR*-*ccoN4* intergenic region, created using data provided by Wurtzel et al. (35). The putative MpaR binding site identified in this study is indicated by an orange rectangle. TSS, transcriptional start site. Bottom: Sequence of the *mpaR*-*ccoN4* intergenic region. Numbers indicate the nucleotide count, running backwards from the ATG. Putative −35 and −10 sites are underlined. (**B**) Expression dynamics of each of the indicated reporter strains during growth in liquid culture. The schematics shown on the right represent the portion of the *mpaR*-*ccoN4* intergenic region included in the reporter construct, with “263 bp” constituting the full intergenic region. A strain containing the full intergenic region in the Δ*mpaR* background is shown as a negative control, and the “137-131 shuffle” strain contains the full intergenic region with bp 137-131 changed to GATATCA. Values shown represent the average of three biological replicates and shading indicates standard deviation. Arrow indicates the onset of stationary phase. (**C**) Promoter regions of the three transcription start sites in the “cyanide-inducible gene cluster” (20, 35). All three contain the same short palindromic repeat 61-62 bp upstream of the TSS (highlighted in orange). Predicted −10 and −35 sites are underlined.

Prior work had identified a GC-rich, putative MpaR-binding motif upstream of *mvfR* (*pqsR*), which codes for a global regulator of quorum sensing (38, 40). The GC-rich sequence at 147 to 138 bp upstream of the *ccoN4* start codon, however, may contribute to but is not required for *ccoN4* expression (**Figure S5**). Identification of the binding motif in the *mvfR* promoter was carried out *in vitro* and with a truncated version of MpaR (38), which could contribute to differences we observed when examining MpaR-dependent P*ccoN4* activity *in vivo*. Though the GC-rich sequence in the *ccoN4* promoter was not required for expression, we instead found that an adjacent palindromic (ATCTGAT) motif was required and therefore might constitute an alternate MpaR binding site.

### Constitutive *mpaR* expression reveals that cyanide is required for full MpaR function

Transcriptomic studies have reported that (i) *mpaR* and its neighboring genes are simultaneously induced by cyanide and (ii) *mpaR*’s neighboring genes are downregulated in a Δ*mpaR* background (20, 38). To further characterize the effects of MpaR and cyanide, we deleted *mpaR* in the Δ*hcn* P*ccoN4-gfp* background and found that the low level of *ccoN4* expression observed in Δ*hcn* was eliminated (**Figure 5A**). The abolishment of basal *ccoN4* expression by *mpaR* deletion in Δ*hcn* hints at the possibility that MpaR has low cyanide-independent activity that is boosted by endogenous cyanide. Together, these observations also suggest two possible scenarios for the regulation of the MpaR/cyanide-controlled gene cluster. In one scenario, cyanide induces production of MpaR and MpaR, in turn, induces transcription of the rest of the gene cluster in a cyanide-independent manner. Alternatively, cyanide might activate MpaR (e.g. by stimulating a conformational change in the protein). To distinguish between these scenarios, we decoupled *mpaR* transcription from MpaR activity by cloning the *mpaR* gene after a synthetic constitutive promoter (41) and inserted this construct at the neutral *glmS* site in a Δ*mpaR* P*ccoN4-gfp* background. Biofilms formed by this MpaR+ P*ccoN4-gfp* strain show high levels of *ccoN4* expression (**Figure 5B**). Removing the capacity to produce cyanide in this strain brought fluorescence down to a level similar to that observed for Δ*hcn* P*ccoN4-gfp* (**Figure 5B**). These results show that cyanide is directly required for full expression of *ccoN4*.

**Figure 5.**
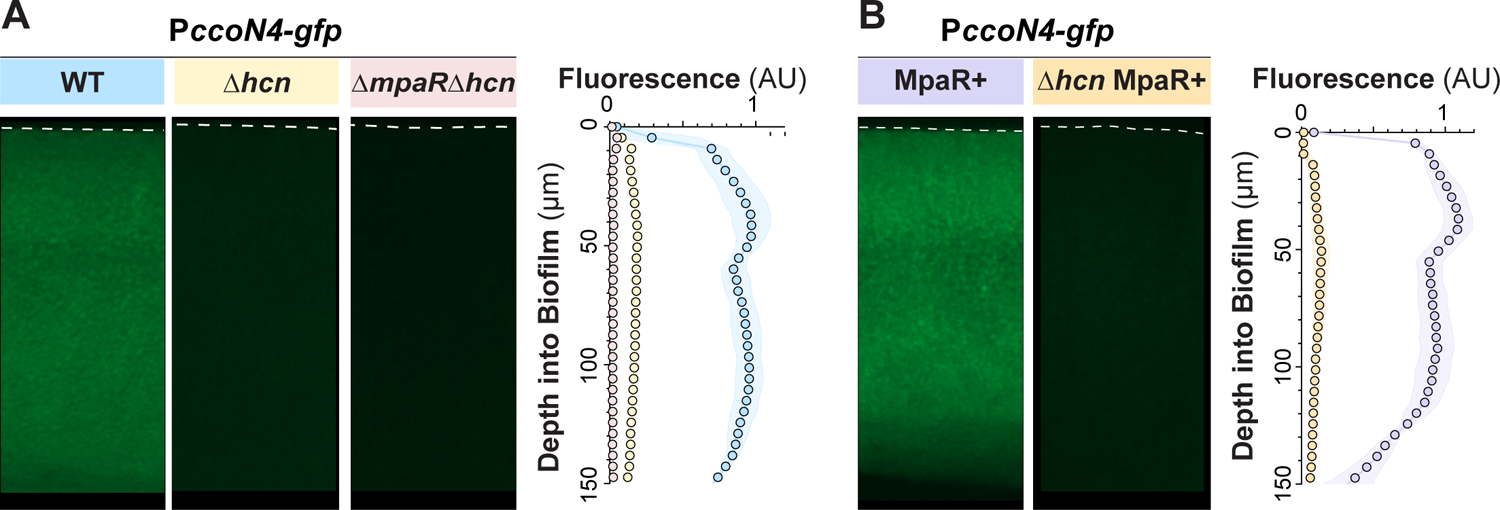
Endogenous cyanide is required for MpaR-driven expression of *ccoN4*. (**A**, **B**) Representative thin-section images of the indicated strains, each containing the P*ccoN4-gfp* reporter construct. Biofilms were grown for 3 days before harvesting and preparation of thin sections. Quantification is shown on the right. Dotted lines indicate the top of each biofilm. For quantification of fluorescence across biofilm depth, experiments were performed with biological triplicates and shading indicates standard deviation.

### Autoregulation of the *PA14_10540-mpaR* operon

Prior transcriptomic analysis carried out by Wurtzel et al. reveals the TSS for *PA14_10540* and indicates that *PA14_10540* and *mpaR* form a transcriptional unit (**Figure 6A**) (35). The region upstream of the *PA14_10540* TSS contains a palindromic ATGNCAT motif (**Figure 4C**), which would suggest control by MpaR (**Figure 4B**) (we note that this experimentally determined TSS is located within the predicted ORF, whose start site may therefore be misannotated in The Pseudomonas Genome Database and BioCyc (42, 43). To test the requirement of MpaR for *PA14_10540* expression, we made a reporter construct (P*PA14_10540-gfp*) containing sequence spanning from 396 bp upstream to 30 bp downstream of the experimentally identified transcription start site (PA14 genomic locus: 908280) (35) (**Figure 6A**). When grown as liquid cultures or biofilms, the P*PA14_10540* reporter strain showed high expression that was eliminated by *mpaR* deletion (**Figure 6B, C**). Examining the effect of the Δ*hcn* mutation on these reporters, we found that the P*PA14_10540* reporter produced an intermediate level of cyanide-independent fluorescence (**Figure 6B, C**). These results indicate that the *PA14_10540*-*mpaR* operon is autoregulated by MpaR and that it is expressed at low levels in the absence of cyanide and at high levels during cyanogenesis.

**Figure 6.**
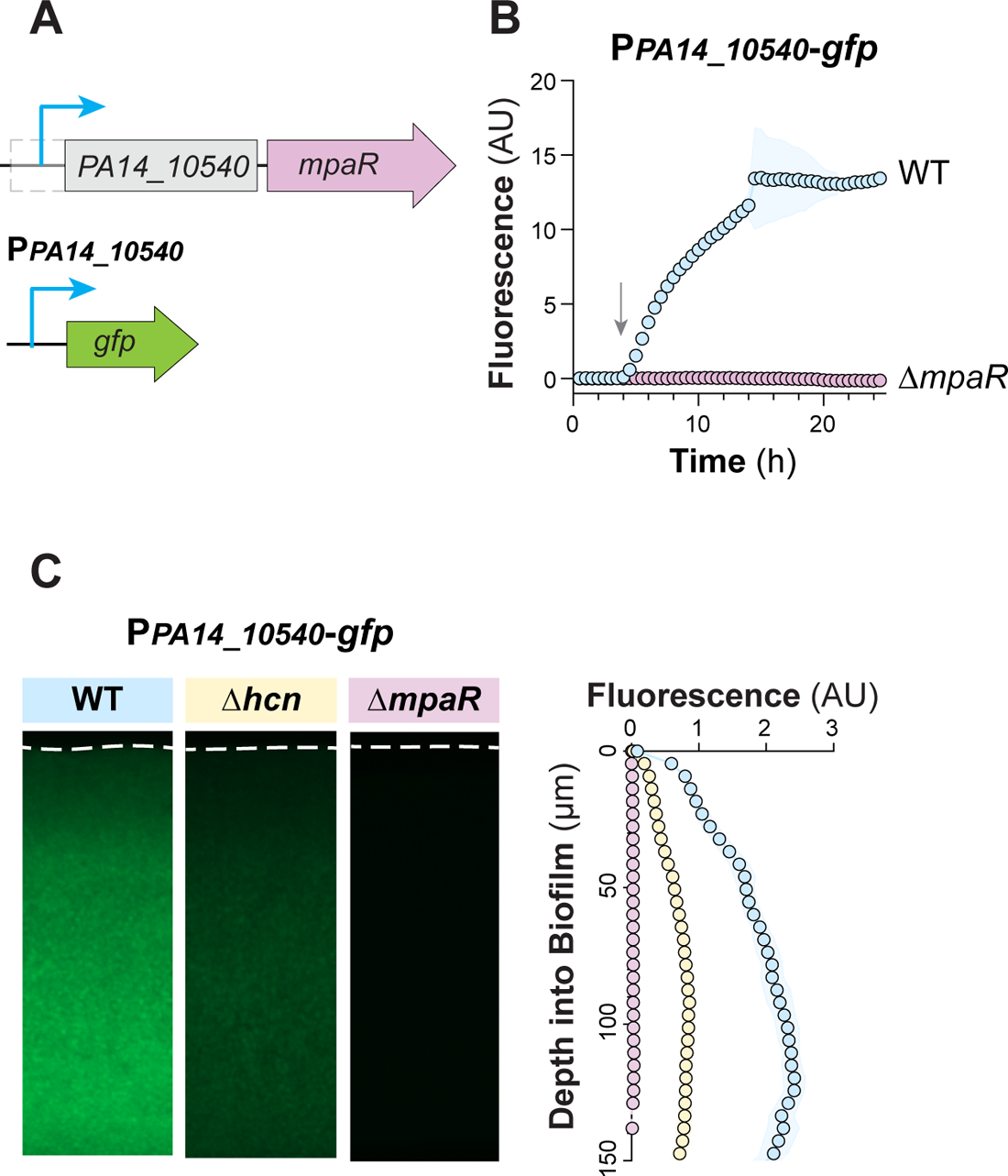
Cyanide-dependent and -independent expression of *mpaR* is driven by the promoter upstream of *PA14_10540*. (**A**) Schematics of the chromosomal region containing *PA14_10540*-*mpaR* and putative transcription start site (additional detail provided in legend for Figure 1A) and of the reporter construct. (**B**) Fluorescence of the indicated strains grown in liquid culture. Data represents 3 biological replicates and shading indicates standard deviation. Arrow indicates the onset of stationary phase. (**C**) Representative thin-section images of the indicated strains, each containing the P*A14_10540-gfp* reporter. Biofilms were grown for 3 days before harvesting and preparation of thin sections. For quantification, shown on the right, experiments were performed with biological triplicates and shading indicates standard deviation.

### Point-mutant phenotypes suggest that MpaR is activated by pyridoxal phosphate

MpaR is a GntR family regulator with an N-terminal, DNA-binding helix-turn-helix (HTH) domain and a C-terminal domain that bears homology to pyridoxal phosphate (PLP)-dependent aminotransferases (**Figure 7A**) (44). We hypothesized that PLP binding could affect MpaR function and examined the structures of PLP-binding proteins that had been co-crystallized with PLP to identify conserved features. PLP-binding pockets contain a conserved lysine residue that covalently binds PLP as a Schiff base and a semi-conserved tyrosine whose ring coordinates the ring of PLP (45, 46). Using an MpaR structure predicted by Alphafold (**Figure 7B**) (47, 48), we identified the corresponding residues in the putative PLP binding site: K314 and Y284 (**Figure 7C,** shown overlaid with solved PLP-bound structure PDB: 1Z2Z). We then created strains in which each of these two residues were changed to alanines, in the P*ccoN4-gfp* reporter background. We found that both point mutations abrogated P*ccoN4-gfp* activity, showing that these two residues are required for MpaR function (**Figure 7D**). MpaR’s PLP-dependent transferase domain is fold type I, and these domains generally bind PLP as homodimers with the active sites brought together (44, 49). Binding of the PLP cofactor may cause MpaR to dimerize in a way that promotes its activity as a transcription factor.

**Figure 7.**
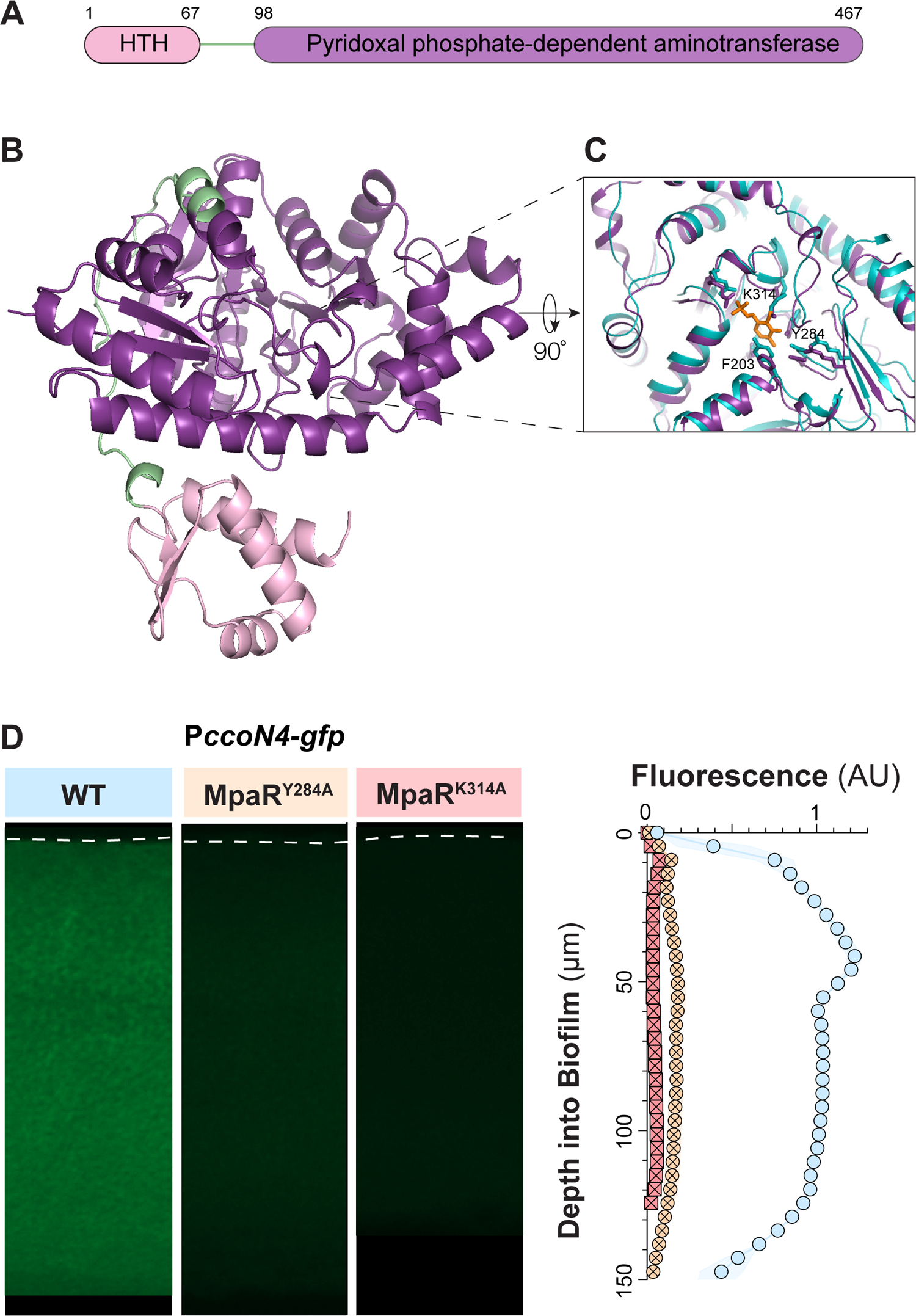
Residues predicted to interact with a pyridoxal phosphate cofactor are required for MpaR-driven expression of *ccoN4*. (**A**) Schematic of the MpaR protein showing putative DNA-binding (HTH) and cofactor (pyridoxal phosphate)-binding domains. Numbers correspond to the amino acid sequence of *P. aeruginosa* PA14 MpaR (44). (**B**) Structure of MpaR generated by Alphafold with domain coloration as shown in (A). (**C**) Reoriented view of MpaR’s (purple) putative cofactor-binding pocket, overlaid on the structure for the putative tRNA pseudouridine synthase D TruD with bound PLP (PDB: 1z2z) (cyan). Tyrosine and lysine residues that extend into the pocket are annotated. (**D**) Representative thin-section images of the indicated strains, each containing the P*ccoN4-gfp* reporter construct. Biofilms were grown for 3 days before harvesting and preparation of thin sections. For quantification, shown on the right, experiments were performed with biological triplicates and shading indicates standard deviation.

### MpaR and CcoN4 mitigate the effects of endogenous cyanide on biofilm development

The responses of *mpaR* and *ccoN4* to endogenous cyanide production in biofilms raised the question of whether these genes play roles in biofilm development. We first tested the *mpaR* deletion for effects on biofilm development using a colony morphology assay (50) We observed that Δ*mpaR* biofilms exhibit a hyperwrinkled phenotype with a distinctive inner ring. In addition, we used thin-section images of biofilms grown on dye-free media to measure biofilm thicknesses and found that Δ*mpaR* biofilms consistently measured approximately 35 µm thinner than their WT counterparts; Δ*ccoN4* biofilms also showed a thinner phenotype, measuring 25 µm thinner than those formed by the WT. Although alterations in phenazine production can affect colony biofilm morphogenesis, HPLC analysis of phenazines produced by Δ*mpaR* biofilms showed subtle changes relative to WT biofilms (**Figure S6**), suggesting that Δ*mpaR* biofilm-development phenotypes arise from other effects. Indeed, deletion of the *hcnABC* operon in the Δ*mpaR* and Δ*ccoN4* backgrounds yielded colony biofilms that more closely resembled WT, and increased the thicknesses of biofilms formed by Δ*mpaR*, Δ*ccoN4*, and even the WT (**Figure 8**). These results indicate that stress arising from the loss of MpaR and CcoN4 can be directly mitigated by removing stress caused by cyanide.

**Figure 8.**
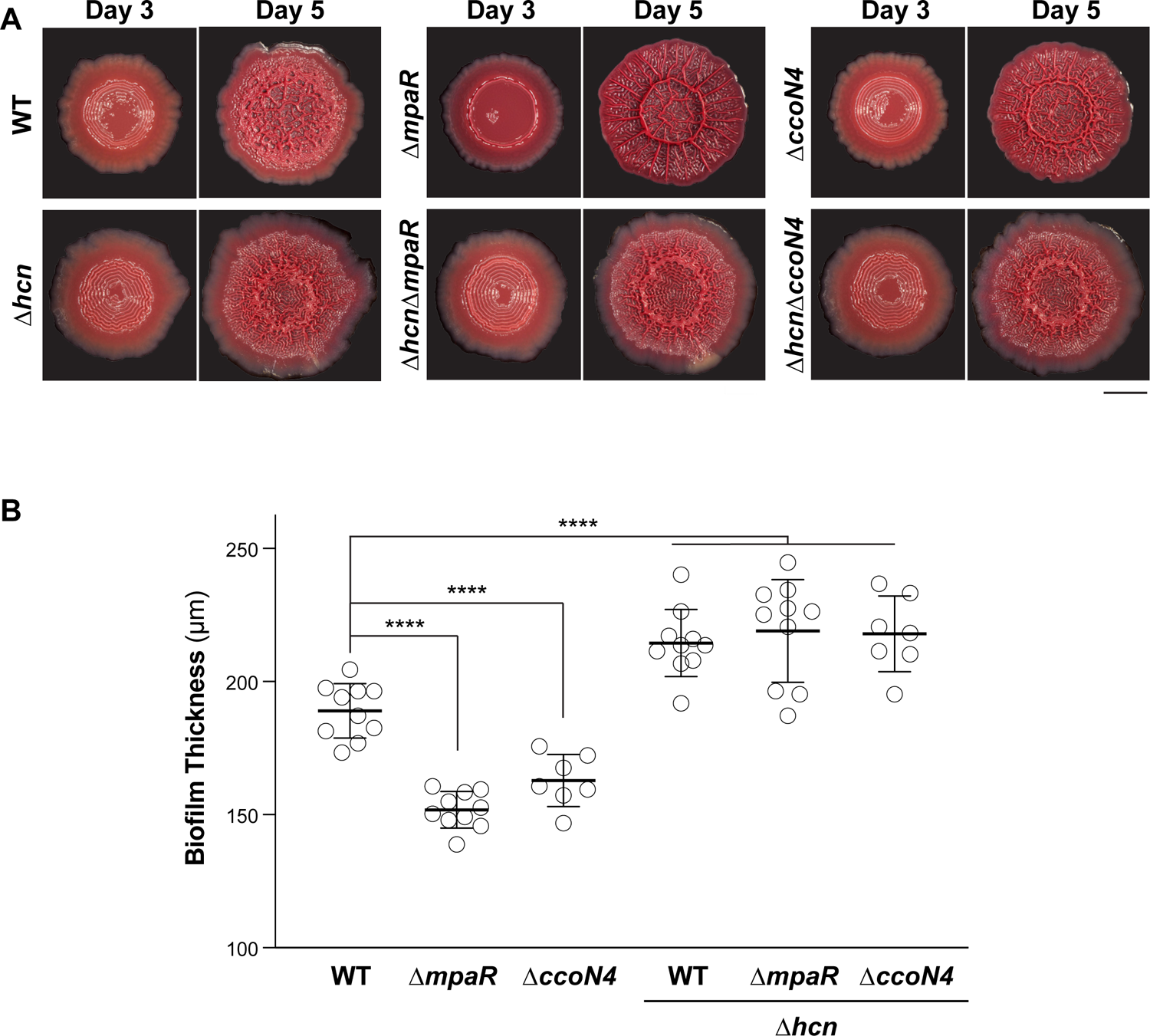
Biofilm development and thickness phenotypes indicate roles for MpaR and CcoN4 in cyanide self-resistance. (**A**) Representative images of biofilms grown in the colony biofilm morphology assay, shown after 3 and 5 days of growth. Scale bar is 5 mm. (**B**) Thicknesses of biofilms formed by the indicated strains. Biofilms were grown for 3 days before harvesting and preparation of thin sections. Data points represent biological replicates and error bars represent standard deviation. P-values were calculated using unpaired, two-tailed t tests (****p<0.0001).

## DISCUSSION

*P. aeruginosa* releases chemically diverse small molecules that have physiological effects on other organisms–including hosts–but also on their producer (51). Among these, cyanide is noteworthy because it is volatile and a potent inhibitor of heme-copper oxidases, which are common components of mitochondrial, bacterial, and archaeal respiratory chains (21, 52, 53). *P. aeruginosa* cyanide production is particularly enigmatic because this bacterium is known for its proficient use of oxygen as a terminal electron acceptor and because it contains six distinct operons encoding subunits of heme-copper oxidases (24, 54). Although *P. aeruginosa* can also produce a cyanide-insensitive (copper-free) oxidase, growth and survival experiments indicate that this oxidase is not necessary for protection against endogenous cyanide under physiological conditions (22, 23). Counterintuitively, *P. aeruginosa* shows strong induction of *ccoN4*, encoding a heme-copper oxidase subunit, when treated with cyanide (20). Experiments with exogenously added cyanide suggest that, even though CcoN4-containing oxidases are predicted to be cyanide-sensitive (19), this subunit acts synergistically with the cyanide-insensitive oxidase to confer cyanide resistance (20).

We set out to investigate the regulation of *ccoN4* expression in the context of *P. aeruginosa* PA14 toxin production and respiration. *ccoN4* lies in a previously identified cluster of six genes that are induced by cyanide (20) and is immediately adjacent to the regulatory gene *mpaR* (38). We confirmed that MpaR is required for *ccoN4* expression, and that full induction of *ccoN4* requires endogenous cyanide production. Furthermore, we identified a short palindromic sequence (ATCTGAT) in the *ccoN4* promoter region that is necessary for this induction. An identical motif appears in the promoter regions of each of the three predicted operons within the so-called “cyanide-inducible gene cluster” (*ccoN4*-*ccoQ4*, *PA14_10540*-*mpaR*, and *nirA-PA14_10560*) at almost the exact same relative location (61, 62, and 62 base pairs upstream of their respective transcription start sites) (35) (**Figure 4C**). Experiments with an engineered strain that constitutively produces MpaR revealed that cyanide production is required for full MpaR-dependent induction of *ccoN4* and not solely for induction of *mpaR*. The fact that the *PA14_10540* promoter contains the putative MpaR binding motif suggests the existence of a positive feedback loop that amplifies the response to cyanide.

Prior transcriptomic studies, comparing WT *P. aeruginosa* to both a cyanide-null mutant (Δ*hcnB*) and a Δ*mpaR* mutant, allow us to make inferences regarding the regulatory effects of cyanide and MpaR. The genes in the “cyanide-inducible gene cluster” stood out among those in the cyanide stimulon due to their high level of induction (20) and also showed high changes in expression in the study comparing the WT and Δ*mpaR* transcriptomes (38) (**Figure S2**). These findings suggest that cyanide affects transcription mainly through MpaR. Consistent with our observation that *ccoN4* expression occurs at low levels in the Δ*hcn* background but is abolished in the Δ*mpaR* background, the MpaR regulon contains genes that are not in the cyanide stimulon–and some of these show strong dependence on MpaR (20, 38)--underscoring the possibility that MpaR exhibits both cyanide-independent and cyanide-dependent activity.

MpaR is predicted to contain an N-terminal helix-turn-helix DNA-binding domain and a C-terminal domain with homology to PLP-dependent aminotransferases, connected by a flexible 31-residue linker (43, 44). These characteristics suggest that MpaR is a member of the MocR subfamily within the GntR family of bacterial transcription factors (55). Although members of the MocR subfamily contain a domain with homology to PLP-dependent aminotransferases, studies have indicated that this domain is not enzymatically active and that–in these proteins–binding of PLP and/or related compounds instead serves as a modulator of transcription factor function (56–59). In *Bacillus clausii* PdxR, for example, it has been shown that the binding of PLP to the C-terminal domain can affect the conformation of the N-terminal domain, thereby changing the protein’s affinity for distinct DNA-binding motifs (60). It should be noted that PLP, a critical cofactor in diverse metabolic pathways, reacts readily with cyanide to generate cyanohydrins (61, 62). This raises the possibility that the effect of cyanide on gene expression is mediated through cellular-level shifts in PLP-associated physiology.

Although organisms that produce cyanide–which include plants, fungi, cyanobacteria, and soil and plant-associated bacteria–risk self-poisoning, they can also benefit from its toxicity to predatory, pathogenic, or competing organisms (52, 63–66). *P. aeruginosa* cyanide production, which can yield concentrations of more than 100 µM (substantially higher than the level needed to kill eukaryotic cells (17, 67)), contributes to its interspecies competitive fitness and its pathogenicity and may also contribute to intraspecies, population-level fitness (68–72). Studies in plants and animals have also highlighted other physiological effects of cyanide production (73), including roles in regulation. Cyanide sensing in plant proteins may occur via chemical modification of cysteine (74); and although other bacterial cyanide-sensing proteins have yet to be identified, a few structures showing the effect of cyanide on ligand binding have been solved (75–77). Our findings suggest a new mechanism of cyanide-dependent control of gene expression in a cyanide-producing, pathogenic bacterium and that could be targeted in therapeutic approaches that weaken *P. aeruginosa’s* self-resistance and toxicity. Moreover, our results point to a possible strategy for the quantification of cyanide that relies on PLP-binding transcription factors, which could be exploited in diverse industrial settings (78–80).

## METHODS

### Strains and Growth Conditions

All bacterial strains used in this study are listed in **Table S1**. For liquid culture growth, single colonies of *Pseudomonas aeruginosa* UCBPP-PA14 (81) were used to inoculate lysogeny broth (LB; 1% tryptone, 1% NaCl, 0.5% yeast extract) (82) or 1% tryptone, as described, and shaken at 37°C, 200 rpm. Generally, liquid precultures were used as inocula for experiments. Biological replicates came from separate clonal source colonies on a streaked agar plate.

### TTC reduction assay

Precultures were grown for 14-16 hours, subcultured 1:100 and grown to an OD 500 nm of ∼0.5. Five µL were spotted onto 60 mL of 1% tryptone, 1% agar [Teknova, A7777] containing 0.004% (w/v) TTC (2,3,5-triphenyl-tetrazolium chloride [Sigma-Aldrich, T8877] in 10 cm X 10 cm X 1.5 cm plates. Plates were incubated for 1 day in the dark at 25°C with >90% humidity and imaged using a flatbed scanner [Epson, E11000XL-GA]. TTC reduction was quantified by measuring the average saturation in the red hue for all pixels in the colony biofilm area.

### Construction of fluorescent reporter strains

All plasmids used in this study are listed in **Table S2**.To construct reporter strains, promoter regions of varying length or with nucleotide changes (as specified in the strain name and in the text) were amplified from the PA14 genome using primers listed in **Table S3** and inserted upstream of the coding sequence of *gfp* using SpeI and XhoI sites in the MCS of the pLD2722 vector. Plasmids were transformed into *E. coli* UQ950 cells and verified by sequencing. Verified plasmids were introduced into PA14 using biparental conjugation with *E. coli* S17-1. Single recombinants were selected on agar plates with M9 minimal medium (47.8 mM Na_2_HPO_4_7H_2_O, 2 mM KH_2_PO_4_, 8.6 mM NaCl, 18.6 mM NH_4_Cl, 1 mM MgSO_4_, 0.1 mM CaCl_2_, 20 mM sodium citrate dihydrate, 1.5% agar) containing 70 µg/mL gentamicin. The plasmid backbone was resolved out of PA14 using Flp-FRT recombination using the pFLP2 plasmid (83) and selection on M9 minimal medium agar plates containing 300 µg/mL carbenicillin. Strains were cured of the pFLP2 plasmid by streaking on LB agar plates without NaCl with 10% w/v sucrose. The presence of *gfp* in final clones was confirmed by PCR.

Shuffled reporters were constructed using an overlap PCR in which two fragments were generated flanking the region to be shuffled with the shuffled sequence introduced with the primers. Then the overlap PCR was performed such that one single fragment was generated from the two fragments. Fragment was inserted into pLD2722 using SpeI and XhoI digest sites and introduced into PA14 as above.

### Construction of deletion mutant strains

All plasmids used in this study are listed in **Table S2.** Markerless deletion mutants in *Pseudomonas aeruginosa* UCBPP-PA14 were made by amplifying ∼1 kb of flanking sequence on each side of the deletion region using the primers listed in **Table S3** and inserting flanks into pMQ30 plasmid through gap repair cloning in *S. cerevisiae* InvSc1 (84). Plasmids were transformed into *E. coli* UQ950, verified by sequencing and introduced into PA14 using biparental conjugation with *E. coli* WM3064. Single recombinants were selected on LB agar plates containing 100 µg/mL gentamicin. Double recombinants were selected on LB without NaCl with 10% w/v sucrose. Deletions were verified by PCR. Combinatorial mutants were made using a single deletion as PA14 parental background. Complements were made as described above by amplifying the flanking regions and the coding sequence together and introduced into the deletion background. Point mutants were created as described for deletion mutants with the exception that the initial primers introduced the specified codon change and included the entire coding region and ∼1 kb of flanking sequence. Constructs were then introduced into the deletion strain and checked for “complementation”.

### Construction of MpaR overexpression strains

The coding region of MpaR was amplified using primers LD4387 and LD4388 and the *rrnB* terminator was amplified from pAKN69 using primers LD4389 and LD4077 such that the two fragments had ∼20 bp overlapping sequence similarity. An overlap PCR was performed using LD4387 and LD4077 and the two fragments (∼75 ng each) as template. The resulting fragment (MpaR coding region with a terminator) was restriction digest cloned into pAKN69 using SphI and NheI restriction enzymes. The resulting plasmid was transformed into *E. coli* WM3064, verified by sequencing, and introduced into PA14 using biparental conjugation with *E. coli* WM3064. Single recombinants were selected on LB agar plates containing 100 µg/mL gentamicin. Plasmid backbones were not resolved out.

### Colony biofilm morphology assay

Precultures grown for 14-16 hours were subcultured 1:100 and grown until OD 500 nm of ∼ 0.5. 10 µL were spotted onto 60 mL of colony morphology medium (1% tryptone, 1% agar [Teknova, A7777] containing 40 µg/mL Congo red dye [VWR, AAAB24310-14] and 20 µg/mL Coomassie blue dye [VWR, EM-3300]) in a 10 cm X 10 cm X 1.5 cm plates. Plates were incubated for up to 5 days at 25°C with >90% humidity and imaged using a flatbed scanner [Epson, E11000XL-GA].

### Biofilm thin sectioning

Sectioning was performed as previously described (85). Briefly, precultures grown for 14-16 hours were subcultured 1:100 and grown until OD 500 nm of ∼ 0.5. Five µL were spotted onto 45 mL/15 mL bilayer 1% tryptone 1% agar [Teknova, A7777] square 10 cm X 10 cm X 1.5 cm plates and incubated for 72 hours at 25°C. Biofilms were overlaid with 15 mL of 1% molten agar, allowed to solidify for 5-10 minutes, then excised by lifting from the bottom layer into histocassettes where they were fixed in 4% paraformaldehyde in PBS (pH 7.2) for 24 hours. Samples were then washed in a series of increasing ethanol concentrations (25%, 50%, 70%, 95% ethanol in PBS, 3 times in 100% ethanol) followed by clearing with 3 washes in Histo-Clear II [National Diagnostics; Fisher Scientific HS-202], then infiltrated with molten paraffin wax [Leica Biosystems, 39601006]. Samples were then embedded in paraffin wax in rectangular wax molds and allowed to solidify for 2-3 hours. Excess wax was removed from the sample blocks and the center of the biofilm was sectioned in 10 µm steps using an automatic microtome [Thermo Fisher Scientific, 905200ER]. Sections were floated in water and mounted on a pre-charged glass slide. Slides air dried overnight then heat fixed at 45°C for 30 minutes. Once cooled, the slides were rehydrated in PBS by reversing the dehydration steps listed above. Sections were mounted with Prolong Diamond Antifade Mountant [Invitrogen; ThermoFisher Scientific, P36965] and covered with coverslips. Coverslips were sealed with quick-dry nail polish and imaged on a ZEISS Axio Zoom.V16. Non-fluorescent parental strains were used as a control for colony auto-fluorescence. In the graphs shown, fluorescent signal has been corrected for the respective parental background fluorescence.

### Liquid culture growth assays

Overnight precultures in biological triplicate were diluted 1:100 into 1% tryptone in 96-well, black-sided, clear-bottomed plate [Greiner Bio-One; Millipore Sigma, M5811] and incubated at 37°C with continuous shaking on the medium setting in a Biotek Synergy H1 plate reader. Growth was assessed by taking OD readings at 500 nm and gfp readings at excitation and emission wavelengths of 480 nm and 510 nm respectively every 30 minutes for 24 hours.

### Cyanide-dependent induction assay

Precultures were grown for 14 to 16 hours, then subcultured 1:100 and grown to an OD 500 nm of ∼ 0.5. Five µL of the cultures were spotted ∼1 cm apart onto 60 mL of 1% tryptone, 1% agar [Teknova, A7777] in a 10 cm X 10 cm X 1.5 cm plates. Plates were incubated for up to 3 days at 25°C with >90% humidity and imaged using a ZEISS Axio Zoom.V16. No additional cyanide was added to the medium.

### High Performance Liquid Chromatography

14-16 hour precultures were subcultured 1:100 and grown until OD 500 nm of ∼ 0.5. 10 µL were spotted onto 4 mL of 1% tryptone, 1% agar [Teknova, A7777] in 30 mm circular petri plates. Plates were incubated for 3 days in the dark at 25°C with >90% humidity. The entire biofilm and agar were harvested by submerging in 5 mL of 100% methanol and rocking overnight at room temperature. The following day, the methanol extracts were filtered through Costar 0.22 µm cellulose acetate Spin-X centrifuge tube filters [Fisher Scientific, 07200386] then 200 µL were loaded into HPLC vials for analysis using high-performance liquid chromatography (Agilent 1100 HPLC system) as previously described (86, 87) and by comparing sample peaks to peaks of pure phenazine standards run as controls. The area under each peak was integrated and used to determine the concentration of each phenazine.

## Acknowledgments

This work was supported by NIH/NIAID grant R01AI103369 to L.E.P.D. We thank Liang Tong for discussion and feedback on the manuscript.

## REFERENCES

1. Evans CR, Kempes CP, Price-Whelan A, Dietrich LEP. 2020. Metabolic Heterogeneity and Cross-Feeding in Bacterial Multicellular Systems. Trends Microbiol 28:732–743.

2. Jo J, Price-Whelan A, Dietrich LEP. 2022. Gradients and consequences of heterogeneity in biofilms. Nat Rev Microbiol 20:593–607.

3. van Gestel J, Vlamakis H, Kolter R. 2015. Division of Labor in Biofilms: the Ecology of Cell Differentiation. Microbiol Spectr 3:MB–0002–2014.

4. Williamson KS, Richards LA, Perez-Osorio AC, Pitts B, McInnerney K, Stewart PS, Franklin MJ. 2012. Heterogeneity in Pseudomonas aeruginosa biofilms includes expression of ribosome hibernation factors in the antibiotic-tolerant subpopulation and hypoxia-induced stress response in the metabolically active population. J Bacteriol 194:2062–2073.

5. Stewart PS, Franklin MJ. 2008. Physiological heterogeneity in biofilms. Nat Rev Microbiol 6:199–210.

6. Armstrong AV, Stewart-Tull DE. 1971. The site of the activity of extracellular products of Pseudomonas aeruginosa in the electron-transport chain in mammalian cell respiration. J Med Microbiol 4:263–270.

7. Hazan R, Que YA, Maura D, Strobel B, Majcherczyk PA, Hopper LR, Wilbur DJ, Hreha TN, Barquera B, Rahme LG. 2016. Auto Poisoning of the Respiratory Chain by a Quorum-Sensing-Regulated Molecule Favors Biofilm Formation and Antibiotic Tolerance. Curr Biol 26:195–206.

8. Voggu L, Schlag S, Biswas R, Rosenstein R, Rausch C, Götz F. 2006. Microevolution of cytochrome bd oxidase in Staphylococci and its implication in resistance to respiratory toxins released by Pseudomonas. J Bacteriol 188:8079–8086.

9. Ciemniecki JA, Newman DK. 2023. NADH dehydrogenases are the predominant phenazine reductases in the electron transport chain of Pseudomonas aeruginosa. Mol Microbiol https://doi.org/10.1111/mmi.15049.

10. Pessi G, Haas D. 2000. Transcriptional control of the hydrogen cyanide biosynthetic genes hcnABC by the anaerobic regulator ANR and the quorum-sensing regulators LasR and RhlR in Pseudomonas aeruginosa. J Bacteriol 182:6940–6949.

11. Sakhtah H, Price-Whelan A, Dietrich LEP. 2013. Regulation of Phenazine Biosynthesis, p. 19–42. In Chincholkar, S, Thomashow, L (eds.), Microbial Phenazines: Biosynthesis, Agriculture and Health. Springer Berlin Heidelberg, Berlin, Heidelberg.

12. Wang Y, Kern SE, Newman DK. 2010. Endogenous phenazine antibiotics promote anaerobic survival of Pseudomonas aeruginosa via extracellular electron transfer. J Bacteriol 192:365–369.

13. Glasser NR, Kern SE, Newman DK. 2014. Phenazine redox cycling enhances anaerobic survival in Pseudomonas aeruginosa by facilitating generation of ATP and a proton-motive force. Mol Microbiol 92:399–412.

14. Jo J, Cortez KL, Cornell WC, Price-Whelan A, Dietrich LE. 2017. An orphan cbb3-type cytochrome oxidase subunit supports Pseudomonas aeruginosa biofilm growth and virulence. Elife 6.

15. Blumer C, Haas D. 2000. Mechanism, regulation, and ecological role of bacterial cyanide biosynthesis. Arch Microbiol 173:170–177.

16. Sanderson K, Wescombe L, Kirov SM, Champion A, Reid DW. 2008. Bacterial cyanogenesis occurs in the cystic fibrosis lung. Eur Respir J 32:329–333.

17. Ryall B, Davies JC, Wilson R, Shoemark A, Williams HD. 2008. Pseudomonas aeruginosa, cyanide accumulation and lung function in CF and non-CF bronchiectasis patients. Eur Respir J 32:740–747.

18. Mitchell R, Brown S, Mitchell P, Rich PR. 1992. Rates of cyanide binding to the catalytic intermediates of mammalian cytochrome c oxidase, and the effects of cytochrome c and poly(L-lysine). Biochim Biophys Acta 1100:40–48.

19. Hirai T, Osamura T, Ishii M, Arai H. 2016. Expression of multiple cbb3 cytochrome c oxidase isoforms by combinations of multiple isosubunits in Pseudomonas aeruginosa. Proc Natl Acad Sci U S A 113:12815–12819.

20. Frangipani E, Pérez-Martínez I, Williams HD, Cherbuin G, Haas D. 2014. A novel cyanide-inducible gene cluster helps protect Pseudomonas aeruginosa from cyanide. Environ Microbiol Rep 6:28–34.

21. Cooper CE, Brown GC. 2008. The inhibition of mitochondrial cytochrome oxidase by the gases carbon monoxide, nitric oxide, hydrogen cyanide and hydrogen sulfide: chemical mechanism and physiological significance. J Bioenerg Biomembr 40:533–539.

22. Zlosnik JEA, Tavankar GR, Bundy JG, Mossialos D, O’Toole R, Williams HD. 2006. Investigation of the physiological relationship between the cyanide-insensitive oxidase and cyanide production in Pseudomonas aeruginosa. Microbiology 152:1407–1415.

23. Arai H, Kawakami T, Osamura T, Hirai T, Sakai Y, Ishii M. 2014. Enzymatic characterization and in vivo function of five terminal oxidases in Pseudomonas aeruginosa. J Bacteriol 196:4206–4215.

24. Kawakami T, Kuroki M, Ishii M, Igarashi Y, Arai H. 2010. Differential expression of multiple terminal oxidases for aerobic respiration in Pseudomonas aeruginosa. Environ Microbiol 12:1399–1412.

25. Alvarez-Ortega C, Harwood CS. 2007. Responses of Pseudomonas aeruginosa to low oxygen indicate that growth in the cystic fibrosis lung is by aerobic respiration. Mol Microbiol 65:153–165.

26. Comolli JC, Donohue TJ. 2004. Differences in two Pseudomonas aeruginosa cbb3 cytochrome oxidases. Mol Microbiol 51:1193–1203.

27. Ducluzeau A-L, Ouchane S, Nitschke W. 2008. The cbb3 oxidases are an ancient innovation of the domain bacteria. Mol Biol Evol 25:1158–1166.

28. de Gier JW, Schepper M, Reijnders WN, van Dyck SJ, Slotboom DJ, Warne A, Saraste M, Krab K, Finel M, Stouthamer AH, van Spanning RJ, van der Oost J. 1996. Structural and functional analysis of aa3-type and cbb3-type cytochrome c oxidases of Paracoccus denitrificans reveals significant differences in proton-pump design. Mol Microbiol 20:1247– 1260.

29. Zufferey R, Preisig O, Hennecke H, Thöny-Meyer L. 1996. Assembly and function of the cytochrome cbb3 oxidase subunits in Bradyrhizobium japonicum. J Biol Chem 271:9114– 9119.

30. Rich PR, Mischis LA, Purton S, Wiskich JT. 2001. The sites of interaction of triphenyltetrazolium chloride with mitochondrial respiratory chains. FEMS Microbiol Lett 202:181–187.

31. Sana TG, Lomas R, Gimenez MR, Laubier A, Soscia C, Chauvet C, Conesa A, Voulhoux R, Ize B, Bleves S. 2019. Differential Modulation of Quorum Sensing Signaling through QslA in Pseudomonas aeruginosa Strains PAO1 and PA14. J Bacteriol 201.

32. Grace A, Sahu R, Owen DR, Dennis VA. 2022. Pseudomonas aeruginosa reference strains PAO1 and PA14: A genomic, phenotypic, and therapeutic review. Front Microbiol 13:1023523.

33. Castric PA. 1975. Hydrogen cyanide, a secondary metabolite of Pseudomonas aeruginosa. Can J Microbiol 21:613–618.

34. Laville J, Blumer C, Von Schroetter C, Gaia V, Défago G, Keel C, Haas D. 1998. Characterization of the hcnABC gene cluster encoding hydrogen cyanide synthase and anaerobic regulation by ANR in the strictly aerobic biocontrol agent Pseudomonas fluorescens CHA0. J Bacteriol 180:3187–3196.

35. Wurtzel O, Yoder-Himes DR, Han K, Dandekar AA, Edelheit S, Greenberg EP, Sorek R, Lory S. 2012. The single-nucleotide resolution transcriptome of Pseudomonas aeruginosa grown in body temperature. PLoS Pathog 8:e1002945.

36. Fenn S, Dubern J-F, Cigana C, De Simone M, Lazenby J, Juhas M, Schwager S, Bianconi I, Döring G, Elmsley J, Eberl L, Williams P, Bragonzi A, Cámara M. 2021. NirA Is an Alternative Nitrite Reductase from Pseudomonas aeruginosa with Potential as an Antivirulence Target. MBio 12.

37. Marckmann D, Trasnea P-I, Schimpf J, Winterstein C, Andrei A, Schmollinger S, Blaby-Haas CE, Friedrich T, Daldal F, Koch H-G. 2019. The cbb3-type cytochrome oxidase assembly factor CcoG is a widely distributed cupric reductase. Proc Natl Acad Sci U S A 116:21166–21175.

38. Wang T, Qi Y, Wang Z, Zhao J, Ji L, Li J, Cai Z, Yang L, Wu M, Liang H. 2020. Coordinated regulation of anthranilate metabolism and bacterial virulence by the GntR family regulator MpaR in Pseudomonas aeruginosa. Mol Microbiol 114:857–869.

39. Trunk K, Benkert B, Quäck N, Münch R, Scheer M, Garbe J, Jänsch L, Trost M, Wehland J, Buer J, Jahn M, Schobert M, Jahn D. 2010. Anaerobic adaptation in Pseudomonas aeruginosa: definition of the Anr and Dnr regulons. Environ Microbiol 12:1719–1733.

40. Déziel E, Gopalan S, Tampakaki AP, Lépine F, Padfield KE, Saucier M, Xiao G, Rahme LG. 2005. The contribution of MvfR to Pseudomonas aeruginosa pathogenesis and quorum sensing circuitry regulation: multiple quorum sensing-regulated genes are modulated without affecting lasRI, rhlRI or the production of N-acyl-L-homoserine lactones. Mol Microbiol 55:998–1014.

41. Lanzer M, Bujard H. 1988. Promoters largely determine the efficiency of repressor action. Proc Natl Acad Sci U S A 85:8973–8977.

42. Karp PD, Billington R, Caspi R, Fulcher CA, Latendresse M, Kothari A, Keseler IM, Krummenacker M, Midford PE, Ong Q, Ong WK, Paley SM, Subhraveti P. 2019. The BioCyc collection of microbial genomes and metabolic pathways. Brief Bioinform 20:1085– 1093.

43. Winsor GL, Griffiths EJ, Lo R, Dhillon BK, Shay JA, Brinkman FSL. 2016. Enhanced annotations and features for comparing thousands of Pseudomonas genomes in the Pseudomonas genome database. Nucleic Acids Res 44:D646–53.

44. Paysan-Lafosse T, Blum M, Chuguransky S, Grego T, Pinto BL, Salazar GA, Bileschi ML, Bork P, Bridge A, Colwell L, Gough J, Haft DH, Letunić I, Marchler-Bauer A, Mi H, Natale DA, Orengo CA, Pandurangan AP, Rivoire C, Sigrist CJA, Sillitoe I, Thanki N, Thomas PD, Tosatto SCE, Wu CH, Bateman A. 2023. InterPro in 2022. Nucleic Acids Res 51:D418– D427.

45. Krupka HI, Huber R, Holt SC, Clausen T. 2000. Crystal structure of cystalysin from Treponema denticola: a pyridoxal 5’-phosphate-dependent protein acting as a haemolytic enzyme. EMBO J 19:3168–3178.

46. Tramonti A, Ghatge MS, Babor JT, Musayev FN, di Salvo ML, Barile A, Colotti G, Giorgi A, Paredes SD, Donkor AK, Al Mughram MH, de Crécy-Lagard V, Safo MK, Contestabile R. 2022. Characterization of the Escherichia coli pyridoxal 5’-phosphate homeostasis protein (YggS): Role of lysine residues in PLP binding and protein stability. Protein Sci 31:e4471.

47. Jumper J, Evans R, Pritzel A, Green T, Figurnov M, Ronneberger O, Tunyasuvunakool K, Bates R, Žídek A, Potapenko A, Bridgland A, Meyer C, Kohl SAA, Ballard AJ, Cowie A, Romera-Paredes B, Nikolov S, Jain R, Adler J, Back T, Petersen S, Reiman D, Clancy E, Zielinski M, Steinegger M, Pacholska M, Berghammer T, Bodenstein S, Silver D, Vinyals O, Senior AW, Kavukcuoglu K, Kohli P, Hassabis D. 2021. Highly accurate protein structure prediction with AlphaFold. Nature 596:583–589.

48. Varadi M, Anyango S, Deshpande M, Nair S, Natassia C, Yordanova G, Yuan D, Stroe O, Wood G, Laydon A, Žídek A, Green T, Tunyasuvunakool K, Petersen S, Jumper J, Clancy E, Green R, Vora A, Lutfi M, Figurnov M, Cowie A, Hobbs N, Kohli P, Kleywegt G, Birney E, Hassabis D, Velankar S. 2022. AlphaFold Protein Structure Database: massively expanding the structural coverage of protein-sequence space with high-accuracy models. Nucleic Acids Res 50:D439–D444.

49. Kirsch JF, Eichele G, Ford GC, Vincent MG, Jansonius JN, Gehring H, Christen P. 1984. Mechanism of action of aspartate aminotransferase proposed on the basis of its spatial structure. J Mol Biol 174:497–525.

50. Friedman L, Kolter R. 2004. Genes involved in matrix formation in Pseudomonas aeruginosa PA14 biofilms. Mol Microbiol 51:675–690.

51. Perry EK, Meirelles LA, Newman DK. 2022. From the soil to the clinic: the impact of microbial secondary metabolites on antibiotic tolerance and resistance. Nat Rev Microbiol 20:129–142.

52. Létoffé S, Wu Y, Darch SE, Beloin C, Whiteley M, Touqui L, Ghigo J-M. 2022. Pseudomonas aeruginosa Production of Hydrogen Cyanide Leads to Airborne Control of Staphylococcus aureus Growth in Biofilm and In Vivo Lung Environments. MBio 13:e0215422.s

53. Pei J, Li W, Kinch LN, Grishin NV. 2014. Conserved evolutionary units in the heme-copper oxidase superfamily revealed by novel homologous protein families. Protein Sci 23:1220– 1234.

54. Williams HD, Zlosnik JEA, Ryall B. 2007. Oxygen, cyanide and energy generation in the cystic fibrosis pathogen Pseudomonas aeruginosa. Adv Microb Physiol 52:1–71.

55. Rigali S, Derouaux A, Giannotta F, Dusart J. 2002. Subdivision of the helix-turn-helix GntR family of bacterial regulators in the FadR, HutC, MocR, and YtrA subfamilies. J Biol Chem 277:12507–12515.

56. Tramonti A, Nardella C, di Salvo ML, Pascarella S, Contestabile R. 2018. The MocR-like transcription factors: pyridoxal 5’-phosphate-dependent regulators of bacterial metabolism. FEBS J 285:3925–3944.

57. Suvorova IA, Rodionov DA. 2016. Comparative genomics of pyridoxal 5’-phosphate-dependent transcription factor regulons in Bacteria. Microb Genom 2:e000047.

58. Edayathumangalam R, Wu R, Garcia R, Wang Y, Wang W, Kreinbring CA, Bach A, Liao J, Stone TA, Terwilliger TC, Hoang QQ, Belitsky BR, Petsko GA, Ringe D, Liu D. 2013. Crystal structure of Bacillus subtilis GabR, an autorepressor and transcriptional activator of gabT. Proc Natl Acad Sci U S A 110:17820–17825.

59. Wu R, Sanishvili R, Belitsky BR, Juncosa JI, Le HV, Lehrer HJS, Farley M, Silverman RB, Petsko GA, Ringe D, Liu D. 2017. PLP and GABA trigger GabR-mediated transcription regulation in Bacillus subtilis via external aldimine formation. Proc Natl Acad Sci U S A 114:3891–3896.

60. Tramonti A, Fiascarelli A, Milano T, di Salvo ML, Nogués I, Pascarella S, Contestabile R. 2015. Molecular mechanism of PdxR – a transcriptional activator involved in the regulation of vitamin B6 biosynthesis in the probiotic bacterium Bacillus clausii. FEBS J 282:2966–2984.

61. Bonavita V. 1960. The reaction of pyridoxal 5-phosphate with cyanide and its analytical use. Arch Biochem Biophys 88:366–372.

62. Naoi M, Ichinose H, Takahashi T, Nagatsu T. 1988. Sensitive assay for determination of pyridoxal-5-phosphate in enzymes using high-performance liquid chromatography after derivatization with cyanide. J Chromatogr 434:209–214.

63. Zdor RE. 2015. Bacterial cyanogenesis: impact on biotic interactions. J Appl Microbiol 118:267–274.

64. Siegień I, Bogatek R. 2006. Cyanide action in plants — from toxic to regulatory. Acta Physiol Plant 28:483–497.

65. Sehrawat A, Sindhu SS, Glick BR. 2022. Hydrogen cyanide production by soil bacteria: Biological control of pests and promotion of plant growth in sustainable agriculture. Pedosphere 32:15–38.

66. Barclay M, Knowles CJ. 2001. Cyanide biodegradation by fungi, p. 335–358. In Fungi in Bioremediation. Cambridge University Press.

67. Zlosnik JEA, Williams HD. 2004. Methods for assaying cyanide in bacterial culture supernatant. Lett Appl Microbiol 38:360–365.

68. Smith P, Cozart J, Lynn BK, Alberts E, Frangipani E, Schuster M. 2019. Bacterial Cheaters Evade Punishment by Cyanide. iScience 19:101–109.

69. Yan H, Asfahl KL, Li N, Sun F, Xiao J, Shen D, Dandekar AA, Wang M. 2019. Conditional quorum-sensing induction of a cyanide-insensitive terminal oxidase stabilizes cooperating populations of Pseudomonas aeruginosa. Nat Commun 10:4999.

70. Nair C, Shoemark A, Chan M, Ollosson S, Dixon M, Hogg C, Alton EWFW, Davies JC, Williams HD. 2014. Cyanide levels found in infected cystic fibrosis sputum inhibit airway ciliary function. Eur Respir J 44:1253–1261.

71. Gallagher LA, Manoil C. 2001. Pseudomonas aeruginosa PAO1 kills Caenorhabditis elegans by cyanide poisoning. J Bacteriol 183:6207–6214.

72. Broderick KE, Chan A, Balasubramanian M, Feala J, Reed SL, Panda M, Sharma VS, Pilz RB, Bigby TD, Boss GR. 2008. Cyanide produced by human isolates of Pseudomonas aeruginosa contributes to lethality in Drosophila melanogaster. J Infect Dis 197:457–464.

73. Randi EB, Zuhra K, Pecze L, Panagaki T, Szabo C. 2021. Physiological concentrations of cyanide stimulate mitochondrial Complex IV and enhance cellular bioenergetics. Proc Natl Acad Sci U S A 118.

74. Gotor C, García I, Aroca Á, Laureano-Marín AM, Arenas-Alfonseca L, Jurado-Flores A, Moreno I, Romero LC. 2019. Signaling by hydrogen sulfide and cyanide through post-translational modification. J Exp Bot 70:4251–4265.

75. Orillard E, Anaya S, Johnson MS, Watts KJ. 2021. Oxygen-Induced Conformational Changes in the PAS-Heme Domain of the Pseudomonas aeruginosa Aer2 Receptor. Biochemistry 60:2610–2622.

76. Fukuyama K, Kunishima N, Amada F, Kubota T, Matsubara H. 1995. Crystal structures of cyanide- and triiodide-bound forms of Arthromyces ramosus peroxidase at different pH values. Perturbations of active site residues and their implication in enzyme catalysis. J Biol Chem 270:21884–21892.

77. Milani M, Ouellet Y, Ouellet H, Guertin M, Boffi A, Antonini G, Bocedi A, Mattu M, Bolognesi M, Ascenzi P. 2004. Cyanide binding to truncated hemoglobins: a crystallographic and kinetic study. Biochemistry 43:5213–5221.

78. Mirajkar AL, Mittapelli LL, Nawale GN, Gore KR. 2018. Synthetic green fluorescent protein (GFP) chromophore analog for rapid, selective and sensitive detection of cyanide in water and in living cells. Sens Actuators B Chem 265:257–263.

79. Dai Z, Boon EM. 2010. Engineering of the heme pocket of an H-NOX domain for direct cyanide detection and quantification. J Am Chem Soc 132:11496–11503.

80. Mondal A, Chattopadhyay SK. 2022. Selective Turn-On Fluorescence Sensing of Cyanide Using the Pyridoxal Platform of a Ni(II) Complex. ACS Omega 7:40941–40949.

81. Rahme LG, Stevens EJ, Wolfort SF, Shao J, Tompkins RG, Ausubel FM. 1995. Common virulence factors for bacterial pathogenicity in plants and animals. Science 268:1899–1902.

82. Bertani G. 2004. Lysogeny at mid-twentieth century: P1, P2, and other experimental systems. J Bacteriol 186:595–600.

83. Hoang TT, Karkhoff-Schweizer RR, Kutchma AJ, Schweizer HP. 1998. A broad-host-range Flp-FRT recombination system for site-specific excision of chromosomally-located DNA sequences: application for isolation of unmarked Pseudomonas aeruginosa mutants. Gene 212:77–86.

84. Shanks RMQ, Caiazza NC, Hinsa SM, Toutain CM, O’Toole GA. 2006. Saccharomyces cerevisiae-based molecular tool kit for manipulation of genes from gram-negative bacteria. Appl Environ Microbiol 72:5027–5036.

85. Cornell WC, Morgan CJ, Koyama L, Sakhtah H, Mansfield JH, Dietrich LEP. 2018. Paraffin Embedding and Thin Sectioning of Microbial Colony Biofilms for Microscopic Analysis. J Vis Exp https://doi.org/10.3791/57196.

86. Sakhtah H, Koyama L, Zhang Y, Morales DK, Fields BL, Price-Whelan A, Hogan DA, Shepard K, Dietrich LEP. 2016. The Pseudomonas aeruginosa efflux pump MexGHI-OpmD transports a natural phenazine that controls gene expression and biofilm development. Proc Natl Acad Sci U S A 113:E3538–47.

87. Dietrich LEP, Price-Whelan A, Petersen A, Whiteley M, Newman DK. 2006. The phenazine pyocyanin is a terminal signalling factor in the quorum sensing network of Pseudomonas aeruginosa. Mol Microbiol 61:1308–1321.

88. Buschmann S, Warkentin E, Xie H, Langer JD, Ermler U, Michel H. 2010. The structure of cbb3 cytochrome oxidase provides insights into proton pumping. Science 329:327–330.

