## Supplementary material for "Cyanide-dependent control of terminal oxidase hybridization by *Pseudomonas aeruginosa* MpaR": SI figures and tables

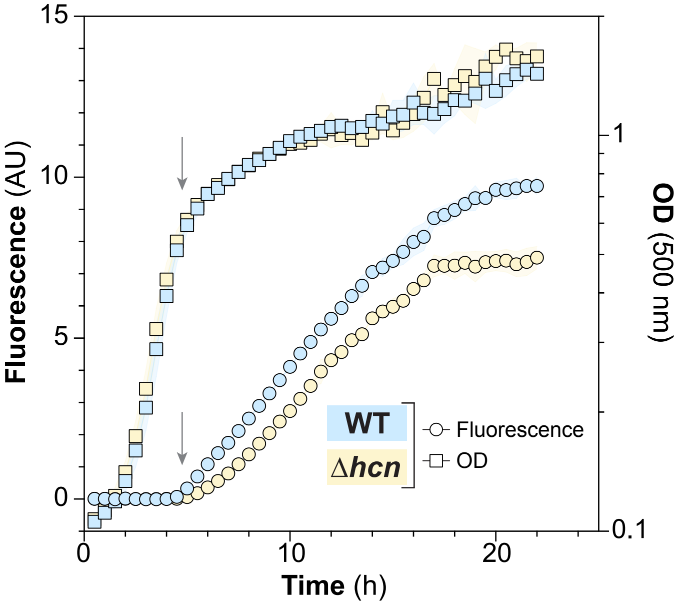


**Figure S1. Endogenous cyanide enhances P*ccoN4-gfp* activity.** Activity of *ccoN4* promoter during growth in liquid culture (1% tryptone) reported as GFP fluorescence. Data represents the average of three biological replicates with shading indicating standard deviation. Arrow indicates the onset of stationary phase.

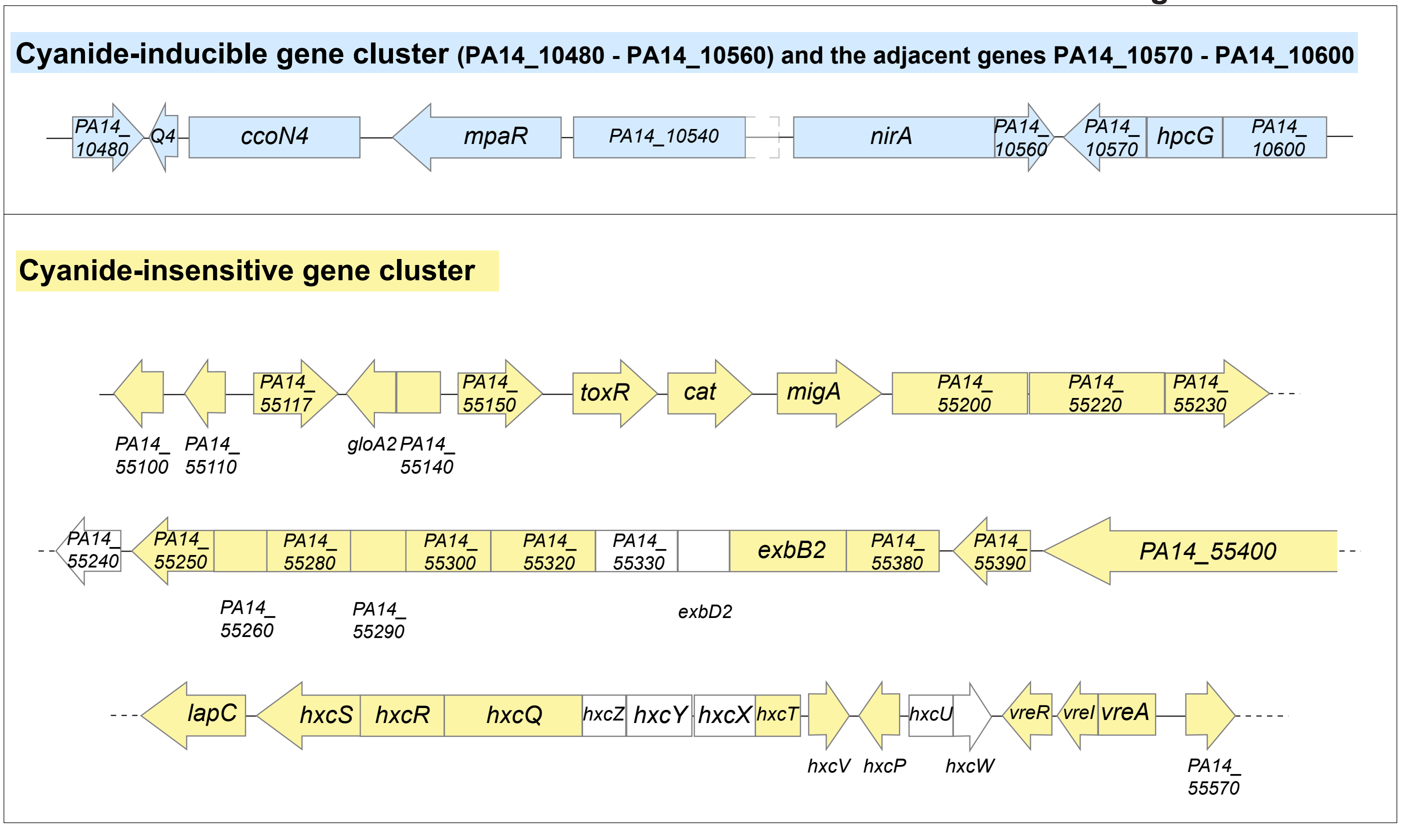


| Gene ID | Gene ID (PA14) | Name | Fold change | P-value | Function |
|---|---|---|---|---|---|
| **PA4132** | PA14_10530 | mpaR | -421.429 | 2.411E-159 | hypothetical protein |
| **PA0713** | PA14_55110 | | -331.686 | 4.013E-112 | hypothetical protein |
| **PA0690** | PA14_55400 | | -107.683 | 1.07E-87 | hypothetical protein |
| **PA4133** | PA14_10500 | ccoN4 | -98.291 | 0E+00 | cytochrome c oxidase subunit (cbb3-type) |
| **PA4130** | PA14_10550 | nirA | -68.082 | 0E+00 | probable sulfite or nitrite reductase |
| **PA4129** | PA14_10560 | | -67.126 | 1.296E-87 | hypothetical protein |
| **PA0689** | - | lapB | -65.337 | 4.851E-53 | low-molecular-weight alkaline phosphatase B |
| **PA0705** | PA14_55180 | migA | -60.714 | 9.662E-55 | alpha-1,6-rhamnosyltransferase MigA |
| **PA0714** | PA14_55100 | | -59.844 | 2.485E-46 | hypothetical protein |
| **PA0704** | PA14_55200 | | -56.050 | 2.459E-47 | probable amidase |
| **PA0706** | PA14_55170 | cat | -43.080 | 4.794E-42 | chloramphenicol acetyltransferase |
| **PA4128** | PA14_10570 | | -41.140 | 7.095E-58 | conserved hypothetical protein |
| **PA0708** | PA14_55150 | | -32.109 | 9.782E-32 | probable transcriptional regulator |
| **PA0709** | PA14_55140 | | -30.483 | 1.099E-29 | hypothetical protein |
| **PA4131** | PA14_10540 | | -25.616 | 0E+00 | probable iron-sulfur protein |
| **PA0688** | PA14_55410 | lapC | -24.192 | 2.614E-26 | probable binding protein component of ABC transporter |
| **PA0703** | PA14_55220 | | -13.842 | 1.126E-15 | probable major facilitator superfamily (MFS) transporter |
| **PA4134** | PA14_10490 | ccoQ4 | -11.920 | 4.72E-92 | hypothetical protein |
| **PA0710** | PA14_55130 | gloA2 | -11.668 | 5.796E-12 | lactoylglutathione lyase |
| **PA0696** | PA14_55320 | | -11.547 | 3.226E-13 | hypothetical protein |
| **PA0673** | PA14_55570 | | -11.112 | 1.227E-12 | hypothetical protein |
| **PA0685** | PA14_55450 | hxcQ | -10.163 | 1.709E-11 | HxcQ probable type II secretion system protein |
| **PA0701a** | PA14_55250 | | -9.550 | 6.089E-11 | hypothetical protein |
| **PA3022** | PA14_24980 | | -8.310 | 1.264E-143 | hypothetical protein |
| **PA0712** | PA14_55117a | | -7.831 | 4.937E-09 | hypothetical protein |

| Gene ID | Gene ID (PA14) | Name | Fold change | P-value | Function |
|---|---|---|---|---|---|
| **PA0679** | PA14_55510 | hxcP | -7.714 | 1.008E-07 | hypothetical protein |
| **PA0692** | PA14_55380 | | -7.513 | 5.386E-08 | hypothetical protein |
| **PA0711** | PA14_55117b | | -7.429 | 1.696E-07 | hypothetical protein |
| **PA0702** | PA14_55230 | | -7.426 | 2.172E-08 | hypothetical protein |
| **PA0676** | PA14_55540 | vreR | -7.166 | 4.816E-08 | sigma factor regulator, VreR |
| **PA4127** | PA14_10590 | hpcG | -7.096 | 4.831E-24 | 2-oxo-hept-3-ene-1,7-dioate hydratase |
| **PA0693** | PA14_55360 | exbB2 | -6.749 | 2.464E-07 | transport protein ExbB2 |
| **PA0701** | PA14_55250 | | -6.688 | 3.966E-07 | probable transcriptional regulator |
| **PA0699** | PA14_55280 | | -6.082 | 9.401E-06 | probable peptidyl-prolyl cis-trans isomerase, PpiC-type |
| **PA0681** | PA14_55490 | hxcT | -6.035 | 5.105E-06 | HxcT pseudopilin |
| **PA0707** | PA14_55160 | toxR | -4.973 | 4.048E-05 | transcriptional regulator ToxR |
| **PA0674** | PA14_55560 | vreA | -4.859 | 1.634E-04 | transcriptional regulators VreA |
| **PA0686** | PA14_55440 | hxcR | -4.453 | 4.338E-04 | HxcR probable type II secretion system protein |
| **PA2331** | PA14_34460 | | -4.283 | 7.562E-50 | hypothetical protein |
| **PA0698** | PA14_55290 | | -4.164 | 1.426E-03 | hypothetical protein |
| **PA3912** | PA14_13300 | | -4.147 | 8.174E-13 | hypothetical protein |
| **PA0687** | PA14_55430 | hxcS | -3.900 | 2.921E-03 | HxcS probable type II secretion system protein |
| **PA0675** | PA14_55550 | vreI | -3.776 | 2.418E-03 | ECF sigma factor, VreI |
| **PA0691** | PA14_55390 | | -3.714 | 4.886E-03 | hypothetical protein |
| **PA4135** | PA14_10480 | | -3.663 | 7.044E-23 | probable transcriptional regulator |
| **PA4126** | PA14_10600 | | -3.531 | 2.242E-11 | probable major facilitator superfamily (MFS) transporter |
| **PA0697** | PA14_55300 | | -3.450 | 7.881E-03 | hypothetical protein |
| **PA0680** | PA14_55500 | hxcV | -3.364 | 1.139E-02 | HxcV putative pseudopilin |
| **PA0700** | PA14_55260 | | -3.308 | 1.226E-02 | hypothetical protein |
| **PA1913** | PA14_39790 | | -3.260 | 4.169E-20 | hypothetical protein |

**Figure S2. Genes whose expression is strongly affected by an *mpaR* deletion are clustered in two genomic regions.** Schematics of the two chromosomal regions containing genes that showed large decreases in expression in a transcriptome study comparing Δ*mpaR* to WT (1). Shaded loci represent those that appeared in the RNAseq.

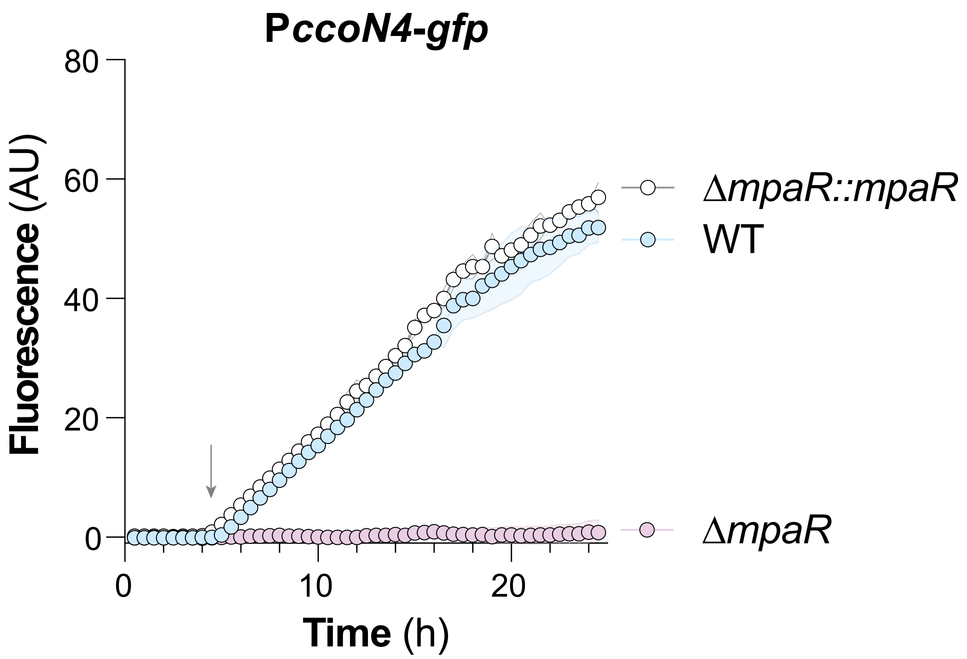


**Figure S3. Complementation of *mpaR* restores expression of *ccoN4*.** Activities of the *ccoN4* promoter during growth in liquid culture (1% tryptone) reported as GFP fluorescence. Data represent the mean of three biological replicates with shading indicating standard deviation. Arrow indicates the onset of stationary phase.

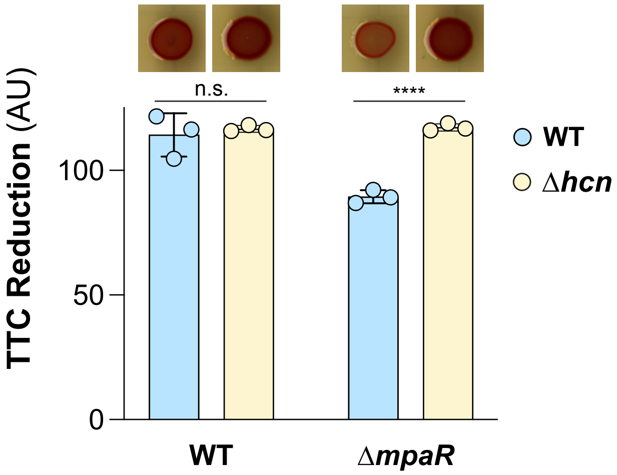


**Figure S4. MpaR is required for full TTC reduction in the presence of cyanide.** (A) TTC reduction (red coloration) by biofilms of the indicated strains, quantified as the average saturation in the red hue for all pixels in the colony area. Data points represent biological replicates and error bars indicate standard deviation.  P-values were calculated using unpaired, two-tailed t tests (****$p<0.0001$).

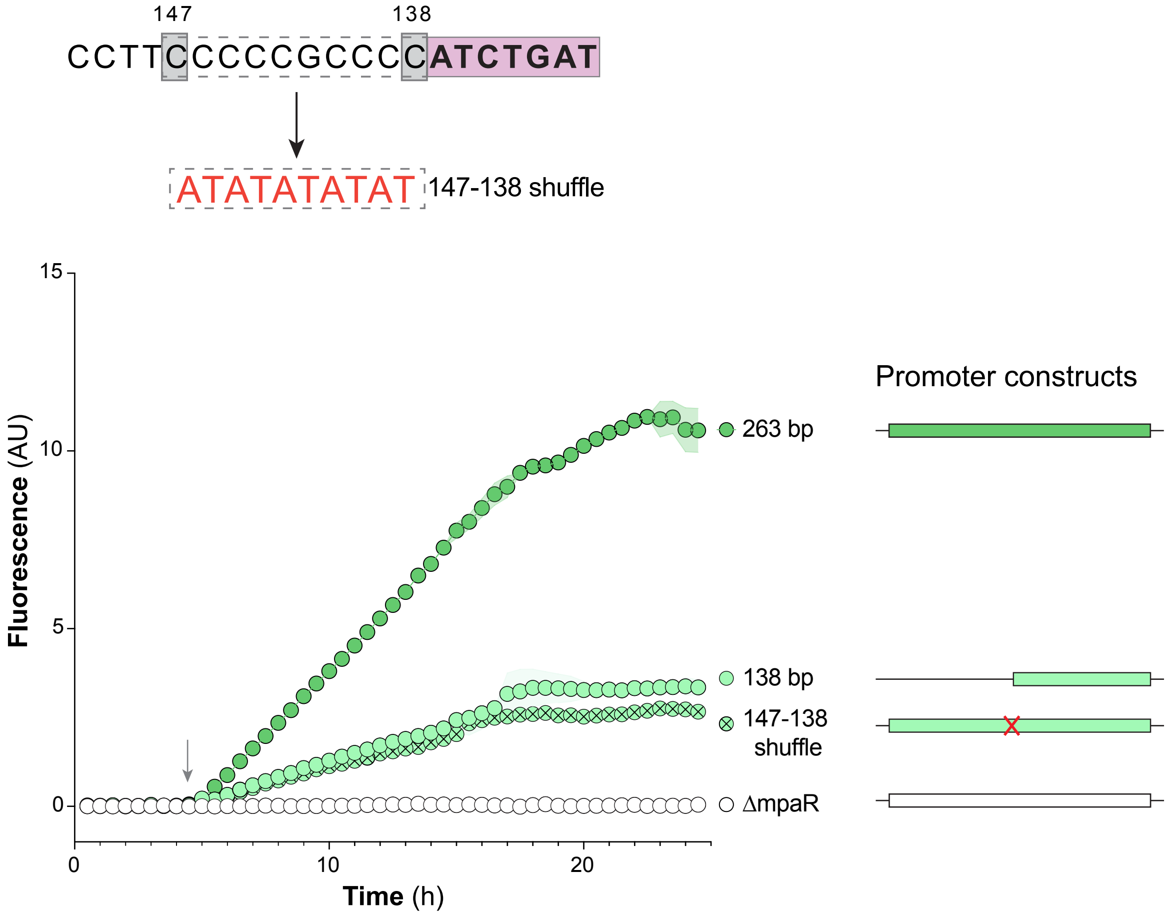


**Figure S5. A GC-rich region upstream of the palindromic motif is required for full expression of the *ccoN4* promoter.** The palindromic motif that is required for *ccoN4* expression (**Figure 4**) is indicated as an orange rectangle. The ten base pairs immediately preceding this motif (147-138) were mutated from CCCCCGCCCC to ATATATATAT in a reporter containing the full intergenic region and the strain is denoted as "147-138 shuffle". A strain containing the full intergenic region in the Δ*mpaR* background is shown as a negative control. Values shown represent the average of three biological replicates and shading indicates standard deviation. Arrow indicates the onset of stationary phase.

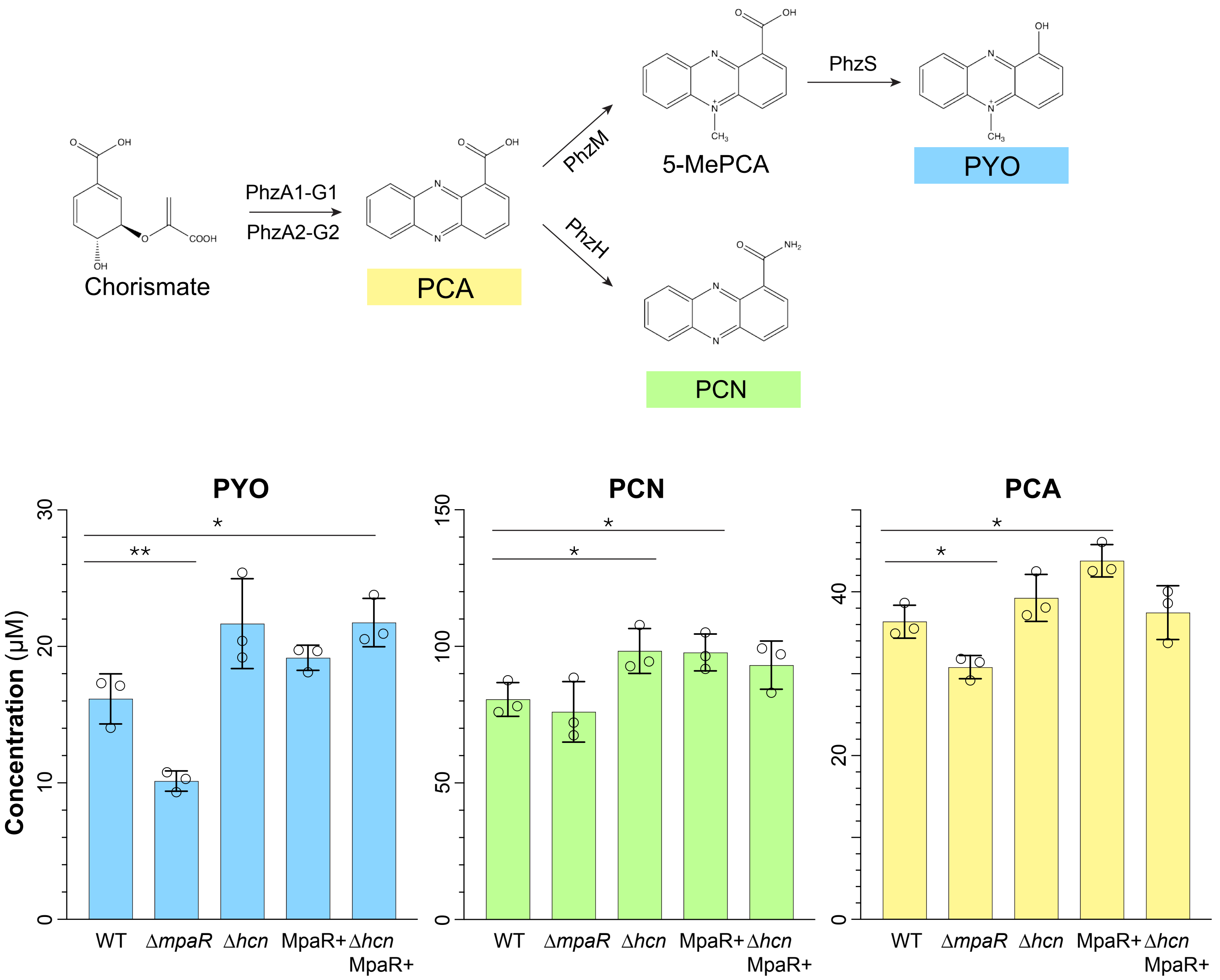


**Figure S6. Phenazine quantification indicates a modest role for MpaR in the regulation of phenazine production.** (**A**) Schematic of the phenazine biosynthetic pathway whereby chorismate is converted to PCA followed by differentiation into derivatives including PCN and PYO. (**B**) High performance liquid chromatography quantification of phenazines produced by biofilms grown for 72 hours on 1% tryptone 1% agar. P-values were calculated using unpaired, two-tailed t tests (*p<0.05; **p<0.01).

### SUPPLEMENTAL TABLES

**Table S1. Bacterial strains used in this study.**

| Number | Strain | Description | Source |
|---|---|---|---|
| *Pseudomonas aeruginosa* strains | | | |
| LD0 | UCBPP-PA14 (WT) | Clinical isolate UCBPP-PA14. | (2) |
| LD2827 | Δ*hcn* | PA14 with the *hcnA-C* (*PA14_36310-PA14_36330*) operon deleted | (3) |
| LD2833 | Δ*ccoN4* | PA14 with *ccoN4* (*PA14_10500*) deleted. | (3) |
| LD2829 | Δ*hcn*Δ*ccoN4* | PA14 with the *hcnA-C* (*PA14_36310-PA14_36330*) operon and *ccoN4* (*PA14_10500*) deleted. | (3) |
| LD1888 | Δ*ccoN1*Δ*ccoN2* | PA14 with *ccoN1* (*PA14_44370*) and *ccoN2* (*PA14_44340*) deleted. | (3) |
| LD2830 | Δ*hcn*Δ*ccoN1*Δ*ccoN2* | PA14 with the *hcnA-C* (*PA14_36310-PA14_36330*) operon, *ccoN1* (*PA14_44370*) and *ccoN2* (*PA14_44340*) deleted. | (3) |
| LD1977 | Δ*ccoN1*Δ*ccoN2*Δ*ccoN4* | PA14 with *ccoN1* (*PA14_44370*), *ccoN2* (*PA14_44340*), and *ccoN4* (*PA14_10500*) deleted. | (3) |
| LD2831 | Δ*hcn*Δ*ccoN1*Δ*ccoN2* Δ*ccoN4* | PA14 with the *hcnA-C* (*PA14_36310-PA14_36330*) operon, *ccoN1* (*PA14_44370*), *ccoN2* (*PA14_44340*), and *ccoN4* (*PA14_10500*) deleted. | (3) |
| LD915 | Δ*anr* | PA14 with *anr* (*PA14_44490*) deleted | (4) |
| LD3884 | Δ*mpaR* | PA14 with *mpaR* (*PA14_10530*) deleted. Made by mating pLD3774 into PA14. | This study |
| LD4867 | Δ*hcn*Δ*mpaR* | PA14 with the *hcnA-C* (*PA14_36310-PA14_36330*) operon and *mpaR* (*PA14_10530*) deleted. Made by mating pLD3774 into LD2827. | This study |
| LD4317 | Δ*mpaR::mpaR* | PA14 Δ*mpaR* (*PA14_10530*) strain with wild-type *mpaR* complemented back into the site of deletion. Made by mating pLD4286 into LD3884. | This study |
| LD2784 | PA14 *attB::*P*ccoN1-gfp* | PA14 containing a construct in the *attB* site | (3) |

| | | | |
|---|---|---|---|
| | | that expresses *gfp* under control of the 500 bp region upstream of *ccoN1* (*PA14_44340*). | |
| LD2786 | PA14 *attB::*P*ccoN2-gfp* | PA14 containing a construct in the *attB* site that expresses *gfp* under control of the 500 bp region upstream of *ccoN2* (*PA14_44370*). | (3) |
| LD4277 | PA14 *attB::*P*ccoN4-gfp* | PA14 containing a construct in the *attB* site that expresses *gfp* under control of the 263 bp region upstream of *ccoN4* (*PA14_10500*). Made by mating pLD4258 into PA14. | This study |
| LD4083 | Δ*anr attB::*P*ccoN1-gfp* | Δ*anr* containing a construct in the *attB* site that expresses *gfp* under control of the 500 bp region upstream of *ccoN1*. Made by mating pLD2766 into LD915. | This study |
| LD4084 | Δ*anr attB::*P*ccoN2-gfp* | Δ*anr* containing a construct in the *attB* site that expresses *gfp* under control of the 500 bp region upstream of *ccoN2*. Made by mating pLD2767 into LD915. | This study |
| LD4342 | Δ*anr attB::*P*ccoN4-gfp* | Δ*anr* containing a construct in the *attB* site that expresses *gfp* under control of the 263 bp region upstream of *ccoN4*. Made by mating pLD4258 into LD915. | This study |
| LD4086 | Δ*mpaR attB::*P*ccoN1-gfp* | Δ*mpaR* containing a construct in the *attB* site that expresses *gfp* under control of the 500 bp region upstream of *ccoN1*. Made by mating pLD2766 into LD3884. | This study |
| LD4087 | Δ*mpaR attB::*P*ccoN2-gfp* | Δ*mpaR* containing a construct in the *attB* site that expresses *gfp* under control of the 500 bp region upstream of *ccoN2*. Made by mating pLD2767 into LD3884. | This study |
| LD4278 | Δ*mpaR attB::*P*ccoN4-gfp* | Δ*mpaR* containing a construct in the *attB* site that expresses *gfp* under control of the 263 bp region upstream of *ccoN4*. Made by mating pLD4258 into LD3884. | This study |
| LD4344 | Δ*hcn attB::*P*ccoN4-gfp* | Δ*hcnA* containing a construct in the *attB* site that expresses *gfp* under control of the 263 bp region upstream of *ccoN4*. Made by mating pLD4258 into LD2827. | This study |
| LD4895 | Δ*hcn*Δ*mpaR attB::*P*ccoN4-gfp* | Δ*hcnA-C* and Δ*mpaR* containing a construct in the *attB* site that expresses *gfp* under control of the 263 bp region upstream of *ccoN4*. Made by mating pLD4258 into | This study |

| | | | |
|---|---|---|---|
| | | LD4867. | |
| LD4318 | Δ*mpaR::mpaR attB::*P*ccoN4-gfp* | Δ*mpaR* containing a construct in the *attB* site that expresses *gfp* under control of the 263 bp region upstream of *ccoN4* with wild-type *mpaR* complemented back into the site of deletion. Made by mating pLD4286 into LD4277. | This study |
| LD4906 | Δ*mpaR* attTn7::P*PA1/04/03-MpaR* | Δ*mpaR* containing a construct in the *attTn7* site that expresses the coding region of *mpaR* under control of the *lac*-derived constitutive *PA1/04/03* promoter. Made by mating pLD4890 into LD3884. | This study |
| LD4907 | Δ*mpaR attTn7::*P*PA1/04/03-MpaR attB::*P*ccoN4-gfp* | Δ*mpaR* containing a construct in the *attTn7* site that expresses the coding region of *mpaR* under control of the *lac*-derived constitutive *PA1/04/03* promoter and a construct in the *attB* site that expresses *gfp* under control of the 263 bp region upstream of *ccoN4*. Made by mating pLD4890 into LD4278. | This study |
| LD4908 | Δ*hcn*Δ*mpaR* attTn7::P*PA1/04/03-MpaR* | Δ*hcnA-C* and Δ*mpaR* containing a construct in the *attTn7* site that expresses the coding region of *mpaR* under control of the *lac*-derived constitutive *PA1/04/03* promoter. Made by mating pLD4890 into LD4867. | This study |
| LD4909 | Δ*hcn*Δ*mpaR attTn7::*P*PA1/04/03-MpaR attB::*P*ccoN4-gfp* | Δ*hcnA-C* and Δ*mpaR* containing a construct in the *attTn7* site that expresses the coding region of *mpaR* under control of the *lac*-derived constitutive *PA1/04/03* promoter and a construct in the *attB* site that expresses *gfp* under control of the 263 bp region upstream of *ccoN4*. Made by mating pLD4890 into LD4895. | This study |
| LD4281 | PA14 *attB::*P*ccoN4(138 bp)-gfp* | PA14 containing a construct in the *attB* site that expresses *gfp* under control of the 138 bp region upstream of *ccoN4*. Made by mating pLD4260 into PA14. | This study |
| LD4336 | PA14 *attB::*P*ccoN4(129bp)-gfp* | PA14 containing a construct in the *attB* site that expresses *gfp* under control of the 129 bp region upstream of *ccoN4*. Made by mating pLD4310 into PA14. | This study |
| LD5000 | PA14 *attB::*P*ccoN4(137bp* | PA14 containing a construct in the *attB* site that expresses *gfp* under control of the 263 | This study |

| | | | |
|---|---|---|---|
| | *shuffle)-gfp* | bp region upstream of *ccoN4* with 7 bp from 137-132 shuffled. Made by mating pLD4975 into PA14. | |
| LD4284 | PA14 *attB*::P*ccoN4(147bp shuffle)-gfp* | PA14 containing a construct in the *attB* site that expresses *gfp* under control of the 263 bp region upstream of *ccoN4* with 10 bp from 147-138 shuffled. Made by mating pLD4263 into PA14. | This study |
| LD5001 | PA14 attB::P*PA14_10540*-gfp | PA14 containing a construct in the *attB* site that expresses *gfp* under control of the ~400 bp surrounding the experimentally identified transcription start site of *PA14_10540*. Made by mating pLD4979 into PA14. | This study |
| LD4996 | Δ*hcn* attB::P*PA14_10540*-gfp | Δ*hcnA-C* containing a construct in the *attB* site that expresses *gfp* under control of the ~400 bp surrounding the experimentally identified transcription start site of *PA14_10540*. Made by mating pLD4979 into LD2827. | This study |
| LD5037 | Δ*mpaR* attB::P*PA14_10540*-gfp | PA14 containing a construct in the *attB* site that expresses *gfp* under control of the ~400 bp surrounding the experimentally identified transcription start site of *PA14_10540*. Made by mating pLD4979 into LD3884. | This study |
| LD5022 | Δ*mpaR*::MpaR$^{Y284A}$ | Y284 of *mpaR* (*PA14_10530*) mutated to alanine in the native locus. Made by mating pLD5006 into LD3884. | This study |
| LD5030 | Δ*mpaR*::MpaR$^{Y284A}$ attB::P*ccoN4-gfp* | Y284 of *mpaR* (*PA14_10530*) mutated to alanine in the native locus. Containing a construct in the *attB* site that expresses *gfp* under the control of the 263 bp region upstream of *ccoN4.* Made by mating pLD4258 into LD5022. | This study |
| LD5018 | Δ*mpaR*::MpaR$^{K314A}$ | K314 of *mpaR* (*PA14_10530*) mutated to alanine in the native locus. Made by mating pLD5007 into LD3884. | This study |
| LD5032 | Δ*mpaR*::MpaR$^{K314A}$ attB::P*ccoN4-gfp* | K314 of *mpaR* (*PA14_10530*) mutated to alanine in the native locus. Containing a construct in the *attB* site that expresses *gfp* under the control of the 263 bp region upstream of *ccoN4.* Made by mating pLD4258 into LD5018. | This study |

| | | | |
|---|---|---|---|
| *E. coli* strains | | | |
| LD44 | UQ950 | E. coli DH5α λ(pir) host for cloning; F-Δ(*argF-lac*)169 Φ80 d*lacZ58*(ΔM15) *glnV44*(AS) *rfbD1 gyrA96*(NalR) *recA1 endA1 spoT1 thi-1 hsdR17 deoR* λpir+ | D. Lies |
| LD661 | BW29427 | Donor strain for conjugation: *thrB1004 pro thi rpsL hsdS lacZ* ΔM15RP4–1360 Δ(*araBAD*)567 Δ*dapA1341*::[*erm pir*(wt)] | W. Metcalf |
| LD69 | β2155 | Helper strain. *thrB1004 pro thi strA hsdsS lacZ*∆M15 (*F' lacZ∆M15 lacI*$^q$ *tra∆36 proA*$^+$ *proB*$^+$) *∆dapA::erm* (Erm$^r$)*pir::*RP4 [*::kan* (Km$^r$) from SM10] | (5) |
| LD2901 | S17-1 | Str$^R$, Tp$^R$, F$^-$ RP4-2-Tc::Mu *aphA*::Tn7 *recA* λpir lysogen | R. Simon |
| *Saccharomyces cerevisiae* strains | | | |
| LD676 | InvSc1 | *MATα/MATα leu2/leu2 trp1-289/trp1-289 ura3-52/ura3-52 his3-Δ1/his3-Δ1* | Invitrogen |

**Table S2. Plasmids used in this study.**

| Plasmid Name | Description | Source |
|---|---|---|
| pMQ30 | Yeast-based allelic-exchange vector; *sacB*$^+$, CEN/ARSH, URA3$^+$, Gm$^R$. | (6) |
| pFLP2 | Site-specific excision vector with cI857-controlled FLP recombinase. encoding sequence, sacB$^+$, Amp$^R$. Used to insert LD2722-based plasmids into P. aeruginosa strains. | (7) |
| pLD2722 | Gm$^R$, Tet$^R$ flanked by Flp recombinase target (FRT) sites to resolve out resistance cassettes. | (3) |
| pAKN69 | Gm$^R$, Cm$^R$ mini-Tn7 $P_{PA1/04/03}$*::yfp* | (8) |
| pLD3774 | ∆*mpaR* (*PA14_10530*) PCR fragment introduced into pMQ30 by gap repair cloning in yeast strain InvSc1. | This study |
| pLD4286 | The CDS of *mpaR* with ~1 kb flanks on either side, introduced into pMQ30 by gap repair cloning in yeast strain InvSc1. | This study |
| pLD2766 | 500 bp upstream of *ccoN1* (*PA14_44340*) PCR fragment in pLD2722. | (3) |

| | | |
|---|---|---|
| pLD2767 | 500 bp upstream of *ccoN2* (*PA14_44370*) PCR fragment in pLD2722. | (3) |
| pLD4258 | 263 bp upstream of *ccoN4* (*PA14_10500*) PCR fragment ligated into pLD2722 using SpeI and XhoI. | This study |
| pLD4260 | 138 bp upstream of *ccoN4* PCR fragment ligated into pLD2722 using SpeI and XhoI. | This study |
| pLD4310 | 129 bp upstream of *ccoN4* PCR fragment ligated into pLD2722 using SpeI and XhoI. | This study |
| pLD4975 | 263 bp upstream of *ccoN4* with the 7 bp between 137 and 132 before the ATG shuffled PCR fragment ligated into pLD2722 using SpeI and XhoI. | This study |
| pLD4263 | 263 bp upstream of *ccoN4* with the 10 bp between 147 and 138 before the ATG shuffled PCR fragment ligated into pLD2722 using SpeI and XhoI. | This study |
| pLD4890 | The CDS of *mpaR* with lambda t0 terminator PCR fragment ligated into pAKN69 using NheI and SphI. | This study |
| pLD4979 | 396 bp upstream to 30 bp downstream of PA14 genomic locus: 908280 PCR fragment ligated into pLD2722 using SpeI and XhoI. | This study |
| pLD5006 | The CDS of *mpaR* with ~1 kb flanks on either side with Y284 changed to an alanine, introduced into pM!30 by gap repair cloning in yeast strain InvSc-1 | This study |
| pLD5007 | The CDS of *mpaR* with ~1 kb flanks on either side with K314 changed to an alanine, introduced into pM!30 by gap repair cloning in yeast strain InvSc-1 | This study |

**Table S3. Primers used in this study.**

| **Primer Number** | **Sequence** |
|---|---|
| Primers for plasmid pLD3774 (used to make ∆*mpaR*) | |
| 3236 | caggcaaattctgttttatcagaccgcttctgcgttctGACCTGGAAAGCCTGTTCTG |
| 3237 | gtctccatcgccttctcctgGTAGGTACGACTGGCGTGC |
| 3238 | ggcacgccagtcgtacctacCAGGAGAAGGCGATGGAG |
| 3239 | ggaattgtgagcggataacaatttcacacaggaaacagctGCCCAGATGTACAGGGTGAT |

| | |
|---|---|
| Primers for plasmid pLD4286 (used to make *mpaR* complement) | |
| 3797 | caggcaaattctgttttatcagaccgcttctgcgttctGACAAGGACACCCTGATCGT |
| 3798 | CAACTCCAGCCAGAGGAAGT |
| 3799 | GCAGTTCCTTCTCCAAGAGC |
| 3800 | ggaattgtgagcggataacaatttcacacaggaaacagctAGGTGGACATGCCGTAGAAC |
| Primers for plasmid pLD4258 (used to make P*ccoN4-gfp*) | |
| 3855 | gattcgactgcactagtCGCCCCGCACACG |
| 3701 | gattcgactgcctcgagCTGTACAGTCCCGAAAGAAATGA |
| Primers for plasmid pLD4260 (used to make P*ccoN4(138bp)-gfp*) | |
| 3860 | gattcgactgcactagtCATCTGATACCCAAATTCCATATC |
| 3701 | gattcgactgcctcgagCTGTACAGTCCCGAAAGAAATGA |
| Primers for plasmid pLD4309 (used to make P*ccoN4(129bp)-gfp*) | |
| 3900 | gattcgactgcactagtCCCAAATTCCATATCGACCTG |
| 3701 | gattcgactgcctcgagCTGTACAGTCCCGAAAGAAATGA |
| Primers for plasmid pLD4975 (used to make P*ccoN4(137bp shuffle)-gfp*) | |
| 3855 | gattcgactgcactagtCGCCCCGCACACG |
| 4491 | CTTCCCCCGCCCCGATATCAACCCAAATTCCATATCGACCT |
| 4492 | AGGTCGATATGGAATTTGGGTTGATATCGGGGCGGGGGAAG |
| 3701 | gattcgactgcctcgagCTGTACAGTCCCGAAAGAAATGA |
| Primers for plasmid pLD4263 (used to make P*ccoN4(147bp shuffle)-gfp*) | |
| 3855 | gattcgactgcactagtCGCCCCGCACACG |
| 3857 | CGATATGGAATTTGGGTATCAGATatatatatatAAGGCGTTCGTGCAGG |
| 3858 | CCTGCACGAACGCCTTatatatatatATCTGATACCCAAATTCCATATCG |
| 3701 | gattcgactgcctcgagCTGTACAGTCCCGAAAGAAATGA |
| Primers for plasmid pLD4890 (used to make P*PA1/04/03-mpaR*) | |
| 4387 | gattcgactgcgcatgctgAAGCGCTACGAGAAATTCG |
| 4388 | actggatctatcaacaggagtccaaTCATCCCGCCAGCG |

| | |
|---|---|
| 4389 | GATCGCCCAGTCGCTGGCGGGATGAttggactcctgttgatagatccag |
| 4077 | acgtacgtacgctagcTTGGATTCTCACCAATAAAAAACGCC |
| Primers for plasmid pLD4979 (used to make P*PA14_10540-gfp*) | |
| 4496 | gattcgactgcactagtGCCTGGTCGTATTCGTCGTA |
| 4497 | gattcgactgcctcgagTTTTCGTAGGCCCATCAGG |
| Primers for plasmid pLD5006 (used to make $mpaR^{Y284A}$) | |
| 4502 | caggcaaattctgttttatcagaccgcttctgcgttctCGATCACCTTCATCGGCTA |
| 4503 | gccgaagtacagttcggcggcCACGTCGTCCTCGATCATC |
| 4504 | gatgatcgaggacgacgtggccGCCGAACTGTACTTCGGC |
| 4505 | ggaattgtgagcggataacaatttcacacaggaaacagctGCCCAGATGTACAGGGTGAT |
| Primers for plasmid pLD5007 (used to make $mpaR^{K314A}$) | |
| 4502 | caggcaaattctgttttatcagaccgcttctgcgttctCGATCACCTTCATCGGCTA |
| 4506 | gtagcccggcgccaggctcgcGGAGAAGGAACTGCAGTGC |
| 4507 | gtgatgcactgcagttccttctccgcgAGCCTGGCGCCGG |
| 4505 | ggaattgtgagcggataacaatttcacacaggaaacagctGCCCAGATGTACAGGGTGAT |

### SUPPLEMENTAL REFERENCES


1. Wang T, Qi Y, Wang Z, Zhao J, Ji L, Li J, Cai Z, Yang L, Wu M, Liang H. 2020. Coordinated regulation of anthranilate metabolism and bacterial virulence by the GntR family regulator MpaR in Pseudomonas aeruginosa. Mol Microbiol 114:857–869.

2. Rahme LG, Stevens EJ, Wolfort SF, Shao J, Tompkins RG, Ausubel FM. 1995. Common virulence factors for bacterial pathogenicity in plants and animals. Science 268:1899–1902.

3. Jo J, Cortez KL, Cornell WC, Price-Whelan A, Dietrich LE. 2017. An orphan cbb3-type cytochrome oxidase subunit supports Pseudomonas aeruginosa biofilm growth and virulence. Elife 6.

4. Lin Y-C, Sekedat MD, Cornell WC, Silva GM, Okegbe C, Price-Whelan A, Vogel C, Dietrich LEP. 2018. Phenazines regulate Nap-dependent denitrification in Pseudomonas aeruginosa biofilms. J Bacteriol https://doi.org/10.1128/JB.00031-18.

5. Dehio C, Meyer M. 1997. Maintenance of broad-host-range incompatibility group P and group Q plasmids and transposition of Tn5 in Bartonella henselae following conjugal plasmid transfer from Escherichia coli. J Bacteriol 179:538–540.

6. Shanks RMQ, Caiazza NC, Hinsa SM, Toutain CM, O'Toole GA. 2006. Saccharomyces cerevisiae-based molecular tool kit for manipulation of genes from gram-negative bacteria. Appl Environ Microbiol 72:5027–5036.

7. Hoang TT, Karkhoff-Schweizer RR, Kutchma AJ, Schweizer HP. 1998. A broad-host-range Flp-FRT recombination system for site-specific excision of chromosomally-located DNA sequences: application for isolation of unmarked Pseudomonas aeruginosa mutants. Gene 212:77–86.

8. Lambertsen L, Sternberg C, Molin S. 2004. Mini-Tn7 transposons for site-specific tagging of bacteria with fluorescent proteins. Environ Microbiol 6:726–732.
